# Critical Environmental Limits: Assessing the Limitations of Core Temperature Inflection Point (CTIP) and Biophysical Modeling

**DOI:** 10.1101/2025.03.21.644619

**Authors:** Faming Wang, Huijuan Xu, Tze-Huan Lei, Yi Xu, Haojian Wang, Lijuan Wang

## Abstract

The Core Temperature Inflection Point (CTIP) method and biophysical modeling are widely used to determine critical environmental limits (CELs), yet their validity under prolonged heat exposure remains unverified. This study evaluated their predictive accuracy by exposing 36 healthy young adults (20 males, 16 females; age: 20.9–22.4 yr) to five counterbalanced 8-hour heat trials in a controlled chamber (36°C/74.5% RH, 40°C/55.0% RH, 44°C/29.2% RH, 47°C/35.6% RH, 50°C/24.5% RH). These conditions were selected based on prior CTIP and biophysical model predictions of CELs. Participants engaged in sedentary office tasks (1.29– 1.67 METs), wore standardized summer clothing (0.39–0.40 clo), and had *ad libitum* access to an electrolyte drink, with a 500-kcal sandwich provided at midday. Rectal temperature (*T*_*rec*_) was continuously monitored. Contrary to model predictions, all five conditions remained compensable (*T*_*rec*_ rise rate ≤ 0.1°C/h), with mean peak *T*_*rec*_ well below heatstroke thresholds (38.2 ± 0.4°C). At 44°C/29.2% RH, females exhibited significantly lower *T*_*rec*_ than males (*p* < 0.05), though steady-state *T*_*rec*_ responses did not differ between sexes (all *p* > 0.10). Collectively, CTIP and biophysical models substantially underestimated CELs, leading to overpredicted heat risk across all trials. These findings challenge the reliability of current predictive methods, suggesting human tolerance may exceed existing estimates. Refining these models is essential for improving heat risk assessment and informing public and occupational health guidelines in a warming climate.

**NEW & NOTEWORTHY:** This study demonstrates that healthy young adults maintained heat balance for 8 hours in five extreme heat conditions (36–50°C), despite the Core Temperature Inflection Point (CTIP) method and biophysical model predictions indicating uncompensable heat stress. These findings challenge the reliability of current assessment tools for prolonged heat exposure and underscore the need for refined models to more accurately predict critical environmental limits, particularly during extreme heatwaves.

## INTRODUCTION

As global temperatures rise due to climate change, extreme heat events are becoming more frequent, intense, and prolonged, posing significant risks to human health and productivity (Mora et al., 2017). Prolonged exposure to extreme heat can lead to heat-related illnesses, ranging from heat exhaustion to life-threatening heat stroke, particularly in vulnerable populations such as the elderly, children, individuals with chronic illnesses, and outdoor workers (Luber and McGeehin, 2008). Understanding human heat tolerance limits is essential for developing effective mitigation strategies (Hanna and Brown, 1983; Sherwood and Ramsay, 2023). To assess these limits, researchers have developed methods to estimate the environmental conditions beyond which humans can no longer maintain thermal balance, often referred to as the uncompensability limit. The most widely used approaches include the Core Temperature Inflection Point (CTIP) method, biophysical modeling, and thermoregulatory modeling (Belding and Kamon, 1973; Kamon and Avellini, 1976; Vecellio et al., 2022; Wolf et al., 2023; Vanos et al., 2023; Xu et al., 2024). While these methods offer rapid estimates, their accuracy under prolonged heat exposure remains uncertain and requires empirical validation (Wang, 2025).

The CTIP method, introduced by Belding and Kamon (1973), identifies the critical environmental limits where heat stress transitions from compensable to uncompensable, marked by a distinct inflection in core temperature as heat gain exceeds heat loss (Avellini et al., 1980). Biophysical modeling, advanced by Vanos et al. (2023), integrates physiological and environmental variables, such as metabolic heat production and evaporative cooling, to estimate heat stress under diverse conditions (Wang et al., 2023). Thermoregulatory modeling (Kang et al., 2019; Xu et al., 2024), exemplified by the six-cylinder thermoregulatory model (SCTM), simulates dynamic physiological responses (e.g., sweating, vasodilation) to estimate critical environmental limits (CELs) for human survivability and livability. Each of these methods aims to refine the theoretical wet-bulb temperature (*T*_*w*_) threshold of 35°C, originally proposed by Sherwood and Huber (2010) as the upper limit for human survival, by incorporating individual variability and real-world factors. However, their validation has largely relied on short-term exposures or simulations, leaving their accuracy for prolonged heat stress unverified.

Prolonged heat exposure, such as an 8-hour workday in hot environments, introduces unique physiological challenges, including cumulative heat storage, hydration deficits, and cardiovascular strain (Kenny et al., 2018; Meade et al., 2024). These factors elevate the risk of heat-related illnesses, yet the CTIP method and biophysical modeling have not been systematically tested under such conditions (Wang, 2025). Previous studies have primarily focused on short-term exposures, limiting the applicability of these models in occupational safety, public health, and climate adaptation planning. Without validation in prolonged heat exposure scenarios, their reliability in predicting human heat tolerance remains uncertain.

In this study, we validated the CTIP method and biophysical modeling against 8-hour heat exposure in a controlled climatic chamber. By comparing their predicted uncompensability limits with observed physiological responses, we assessed their reliability and accuracy under prolonged heat stress. Specifically, we examined core temperature trajectories, hydration status, and metabolic heat production to determine how well these models predict human thermal limits over extended exposure periods. Our findings will clarify the strengths and limitations of these methods, enhancing their applicability for real-world heat stress predictions and informing strategies for mitigating heat-related health risks in a warming world.

## METHODS

### Ethical approval and human participants

This study was approved by the Institutional Review Board of Xi’an University of Science and Technology (approval number: XUST-IRB225003) and registered in the Chinese Clinical Trial Registry (ChiCTR) (registration number: ChiCTR2300071853). All participants provided verbal and written informed consent prior to enrollment, in accordance with the Declaration of Helsinki.

Thirty-six healthy young adults participated in the study: 20 males (age: 22.4 ± 0.9 years, body surface area: 1.83 ± 0.11 m^2^, body mass index [BMI]: 22.3 ± 2.1 kg/m^2^) and 16 females (age: 20.9 ± 2.0 years, body surface area: 1.60 ± 0.12 m^2^, BMI: 21.1 ± 2.2 kg/m^2^). Female participants were tested during the early follicular phase of their menstrual cycle (5 ± 3 days), a period characterized by low estrogen and progesterone levels (Bouman et al., 2005). Participants had low-to-moderate physical activity levels: males exercised 4.9 ± 2.0 times per week (jogging, badminton, strength training), while females exercised 3.0 ± 1.5 times per week (jogging, tennis, basketball, badminton). To ensure a representative sample of young, physically active adults, all participants were screened for cardiovascular, respiratory, and metabolic disorders and had no history of heat-related illnesses or medications affecting thermoregulation.

### Test conditions

To validate the Core Temperature Inflection Point (CTIP) method and biophysical modeling under prolonged heat exposure, environmental conditions were selected based on prior predictions of uncompensability limits. The CTIP method, as defined by Vecellio et al. (2022), established critical relative humidity (RH) thresholds at various dry-bulb temperatures: 74.0% RH at 36°C (wet-bulb temperature, *T*_*W*_=31.9°C), 64.5% RH at 38°C (*T*_*W*_=32.0°C), 53.5% RH at 40°C (*T*_*W*_=31.6°C), 29.2% RH at 44°C (*T*_*W*_=28.7°C), 21.0% RH at 47.3°C (*T*_*W*_=28.1°C), and 13.2% RH at 50.6°C (*T*_*W*_=26.5°C). For validation, these conditions were selected, representing moderate to extreme heat stress: 36°C with 74.5% RH (*T*_*W*_=32.0°C), 40°C with 55.0% RH (*T*_*W*_=32.0°C), and 44°C with 29.2% RH (*T*_*W*_=28.7°C). Biophysical modeling predictions by Vanos et al. (2023) identified survivability limits across various RH values, ranging from 88.9% at 35°C to 16.1% at 52°C dry-bulb temperature. Two conditions were selected to test these predictions: 47°C with 35.6% RH (*T*_*W*_=33.0°C) and 50°C with 24.5% RH (*T*_*W*_=31.4°C). At these conditions, Vanos et al. (2023) estimated that young adults (18–40 years) would reach a core temperature of 43.0°C within 6 hours, indicative of heatstroke risk (Pal and Eltahir, 2016). These five conditions (Figure 1) were implemented in a controlled climatic chamber to assess the reliability of CTIP and biophysical modeling predictions over an 8-hour exposure period. The air velocity in the chamber was maintained at 0.20 ± 0.05 m/s, representing a static environment. The CO_2_ concentration was controlled at 947 ± 32 ppm to ensure optimal air quality during the trials (Satish et al., 2012).

**Figure 1.**
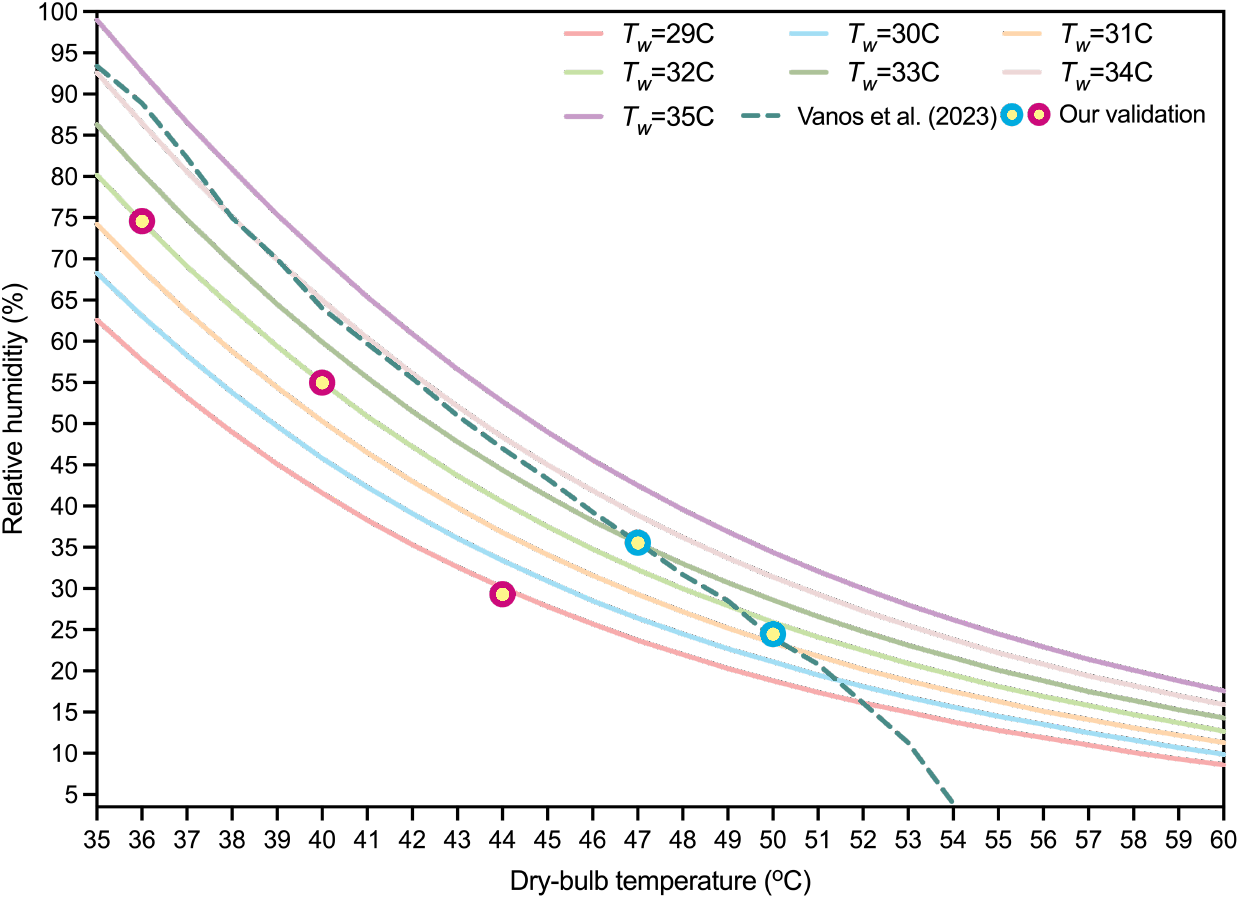
Environment conditions for validating the CTIP method and biophysical modeling. The five validation conditions covered wet-bulb temperatures (*T*_*w*_) ranging from 28.4°C to 33.0°C. The metabolic rates reported in the CTIP study by Vecellio et al. (2022) and the biophysical modeling simulations by Vanos et al. (2023) were 1.3-1.5 METs and 1.5 METs, respectively. Filled circles with a pink border and yellow interior represent the three conditions selected for validating the CTIP method, while filled circles with a cyan border and yellow interior denote the two conditions chosen for validating the biophysical modelling. Additionally, clothing insulation in the CTIP method by Vecellio et al. (2022) and the biophysical modeling simulations by Vanos et al. (2023) was approximately 0.36 clo.

### Protocol overview

Participants completed five counterbalanced 8-hour heat exposure trials, each simulating one of the specified environmental conditions outlined above (36°C/74.5% RH, 40°C/55.0% RH, 44°C/29.2% RH, 47°C/35.6% RH, 50°C/24.5% RH). Trials were conducted in a controlled environmental chamber (Espec Corp., Osaka, Japan) between November 2024 and March 2025 to minimize the seasonal heat acclimatization effects on core temperature fluctuations (Buono et al., 1998). Each trial took place from 9:00 a.m. to 5:00 p.m., with a 5-day washout period between sessions to allow full physiological recovery.

Before each trial, participants provided urine samples to verify adequate hydration, defined as a urine specific gravity (USG) below 1.020 (Oppliger et al., 2005). If USG exceeded this threshold, participants consumed an isotonic electrolyte drink (Nuotelande: 10.8 mg zinc, 0.5 g carbohydrates, 283 mg sodium; Jinan, China) until compliance was achieved. Pre-trial measurements included nude body weight and venous blood sampling. During trials, participants were seated in office chairs, engaging in sedentary office tasks (e.g., reading, computer work). They wore standardized clothing, including short-sleeve T-shirts, sports bras (for females), underwear, trousers, socks, and sports shoes, yielding insulation values of 0.39 clo (males) and 0.40 clo (females) (ASHRAE, 2023).

### Hydration and dietary standardization

Participants had *ad libitum* access to an electrolyte drink throughout the 8-hour heat exposure to maintain hydration, drinking as needed to prevent thirst rather than following a fixed intake schedule. To avoid potential cooling effects, the beverage was stored at 37°C, matching body temperature. Restroom breaks were permitted as needed to maintain comfort and normal fluid regulation.

Dietary intake was standardized across all trials, with participants consuming a 500-kcal (2092 kJ) sandwich at 12:00 p.m. This controlled energy intake minimized dietary influences on metabolic rate and physiological responses (Lei et al., 2025).

### Measurements

#### Body temperature

Core temperature (*T*_rec_) was continuously monitored using a rectal thermistor (YSI401, Yellow Spring Instrument, Yellow Springs, OH; accurate: ±0.1°C) inserted 10 cm beyond the anal sphincter. Rectal thermometry was chosen as an index of deep body temperature due to its resistance to fluid intake effects, ensuring accurate measurements even as participants consumed the standardize meal during the 8-hour heat exposures. Mean skin temperature (*T*_sk_) was calculated using four iButton dataloggers (DS1992L, Maxim Integrated Product Inc., San Jose, CA) placed at the chest, upper arm, thigh, and calf. *T*_sk_ was determined using Ramanathan’s formula ( *T*_*sk*_ = 0.3 ×(*T*_*chest*_ + *T*_*upper arm*_) + 0.2 × (*T*_*thigh*_ + *T*_*calf*_ ) (Ramanathan, 1964).

#### Metabolic rate

Metabolic rate was measured at two time points: at the start (15 minutes after exposure onset) and at the end of the 8-hour exposure, using a COSMED K5 wearable metabolic system (COSMED srl, Rome, Italy). Participants remained seated and engaged in sedentary office tasks (e.g., reading, computer work) during measurements. Each measurement lasted 8 minutes, with the final 5 minutes used for analysis to ensure data stability. To maintain accuracy, the COSMED K5 system was calibrated daily using high-grade calibration gases (GASCO: 5% CO_2_, 21% O_2_, balance N_2_; Airgas Therapeutics LLC, Oldsmar, FL) and a 3-liter calibration syringe (Hans Rudolph 3L Calibration Syringe), following the manufacturer’s instructions. Metabolic rates were expressed in metabolic equivalents (METs) by normalizing oxygen consumption (mL·kg^−1^·min^−1^) to a standard value of 3.5 mL·kg^-1^·min^-1^ (Jetté et al., 1990).

#### Hydration status

Hydration status was monitored throughout the exposure. Participants were instructed to drink *ad libitum*, ensuring they avoided thirst sensation. Urine samples were collected both pre- and post-exposure to assess hydration levels. Urine specific gravity (USG) was measured using a refractometer (Atago PAL-10S, cat. No. 4410, Atago Co. Ltd., Tokyo, Japan; accuracy: ±0.001; range: 1.000 to 1.060).

#### Data processing and statistical analysis

Individual rectal temperature data were smoothed using a spline function in R software (version 4). The rate of core temperature change was calculated every 5 minutes. Steady state temperature was defined by one of two criteria: a core temperature change rate of ≤ 0.10°C/h after 3 hours of exposure or a core temperature fluctuation of ≤ 0.1°C during the final 30 minutes of exposure. If the second criterion was met, the steady-state timing point was set at the 450th minute.

Statistical analyses were performed using IBM SPSS Statistics (version 20, IBM Corp., Armonk, NY), and figures were generated in GraphPad Prism (version 10.3.1, GraphPad Software LLC, Boston, MA). Data normality was assessed via the Kolmogorov-Smirnov test, and variance homogeneity via Levene’s test. A one-way mixed-model analysis of variance (ANOVA) examined the effects of time (9:00 a.m.–5:00 p.m.) on *T*_*rec*_, with sex as a between-subjects factor. Bonferroni-corrected post hoc tests were conducted following significant main or interaction effects. Data are presented as means ± standard deviation (SD), with statistical significance set at *p* ≤ 0.05.

## RESULTS

### Hydration status

Mean urine specific gravity (USG) values remained within the euhydrated range (≤ 1.020, see Table S1 in the **Supplemental Material**) across all five experimental conditions, indicating that participants maintained adequate hydration throughout the 8-hour heat exposure. While minor fluctuations in USG were observed between pre- and post-exposure measurements, these variations remained within normal limits, suggesting that fluid balance was effectively regulated. No evidence of dehydration was detected in either males or females at any tested condition.

### Environmental conditions inside the chamber

The actual environmental conditions recorded for validating the CTIP method were 36.1 ± 0.1°C with 74.1 ± 0.2% RH, 39.9 ± 0.3°C with 55.7 ± 0.7% RH, and 44.0 ± 0.1°C with 29.2 ± 0.2% RH, corresponding to wet-bulb temperatures of 32.0°C, 32.0°C and 28.7°C, respectively. Similarly, the two conditions chosen for validating the biophysical modeling were 47.1 ± 0.1°C with 35.8 ± 0.5% RH and 50.1 ± 0.2°C with 24.0 ± 0.3% RH, corresponding to wet-bulb temperatures of 33.1°C and 31.3°C.

### Metabolic rate

The metabolic rate increased significantly by the end of the 8-hour heat exposure across all environmental conditions (*p* < 0.05; Table 1). The rise in MET from pre-exposure to post-exposure ranged from 15.4% to 20.1% in males and 17.8% to 20.3% in females, depending on the environmental conditions. Notably, males had consistently higher MET values than females throughout the trials (*p* < 0.01), with sex differences being statistically significant at both time points in all conditions (*p* < 0.05). The largest increase in MET was observed in the 50°C, 24.5% RH condition for males (20.1% increase) and the 47°C, 35.6% RH condition for females (20.3% increase), highlighting a trend where both sexes experienced greater metabolic elevations in extreme heat conditions.

**Table 1.**
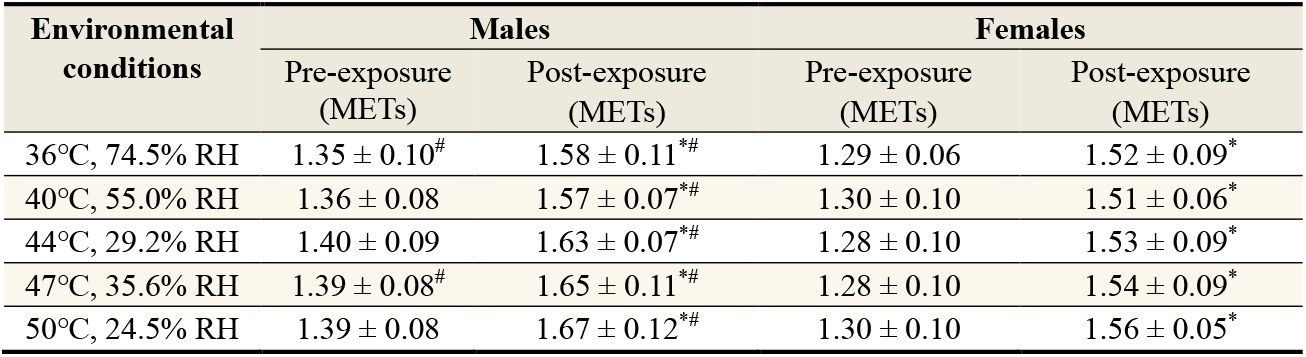
Metabolic equivalent (MET) before and at the end of the exposure for both sexes (males: n=20; females: n=16). Data were analyzed using one-way repeated ANOVA. *, indicates a significant difference from Pre-exposure (*p* < 0.05); ^#^, indicates a significant difference between sexes (*p* < 0.05).

| Environmental conditions | Males |  | Females |  |
| --- | --- | --- | --- | --- |
|  | Pre-exposure (METs) | Post-exposure (METs) | Pre-exposure (METs) | Post-exposure (METs) |
| 36°C, 74.5% RH | 1.35 ± 0.10<sup>#</sup> | 1.58 ± 0.11<sup>*#</sup> | 1.29 ± 0.06 | 1.52 ± 0.09<sup>*</sup> |
| 40°C, 55.0% RH | 1.36 ± 0.08 | 1.57 ± 0.07<sup>*#</sup> | 1.30 ± 0.10 | 1.51 ± 0.06<sup>*</sup> |
| 44°C, 29.2% RH | 1.40 ± 0.09 | 1.63 ± 0.07<sup>*#</sup> | 1.28 ± 0.10 | 1.53 ± 0.09<sup>*</sup> |
| 47°C, 35.6% RH | 1.39 ± 0.08<sup>#</sup> | 1.65 ± 0.11<sup>*#</sup> | 1.28 ± 0.10 | 1.54 ± 0.09<sup>*</sup> |
| 50°C, 24.5% RH | 1.39 ± 0.08 | 1.67 ± 0.12<sup>*#</sup> | 1.30 ± 0.10 | 1.56 ± 0.05<sup>*</sup> |

### Body temperatures

All five experimental conditions represented compensable heat stress for both sexes (Figure 2). Among male participants, the time to reach thermoregulatory steady state across the five conditions (36°C/74.5% RH, 40°C/55.0% RH, 44°C/29.2% RH, 47°C/35.6% RH, 50°C/24.5% RH) was 278 ± 61, 288 ± 100, 216 ± 38, 291 ± 89, and 260 ± 64 minutes, respectively. For females, the corresponding times were 262 ± 48, 256 ± 33, 221 ± 56, 249 ± 70, and 227 ± 50 minutes. Core temperatures during the 8-hour exposure were generally comparable between sexes, except in the 44°C/29.2% RH condition (Figure 2C, *p* < 0.01). No significant sex differences in core temperature were observed in other conditions (all *p* > 0.10). Individual core temperature responses and rates of core temperature change are provided in the **Supplemental Material**.

**Figure 2.**
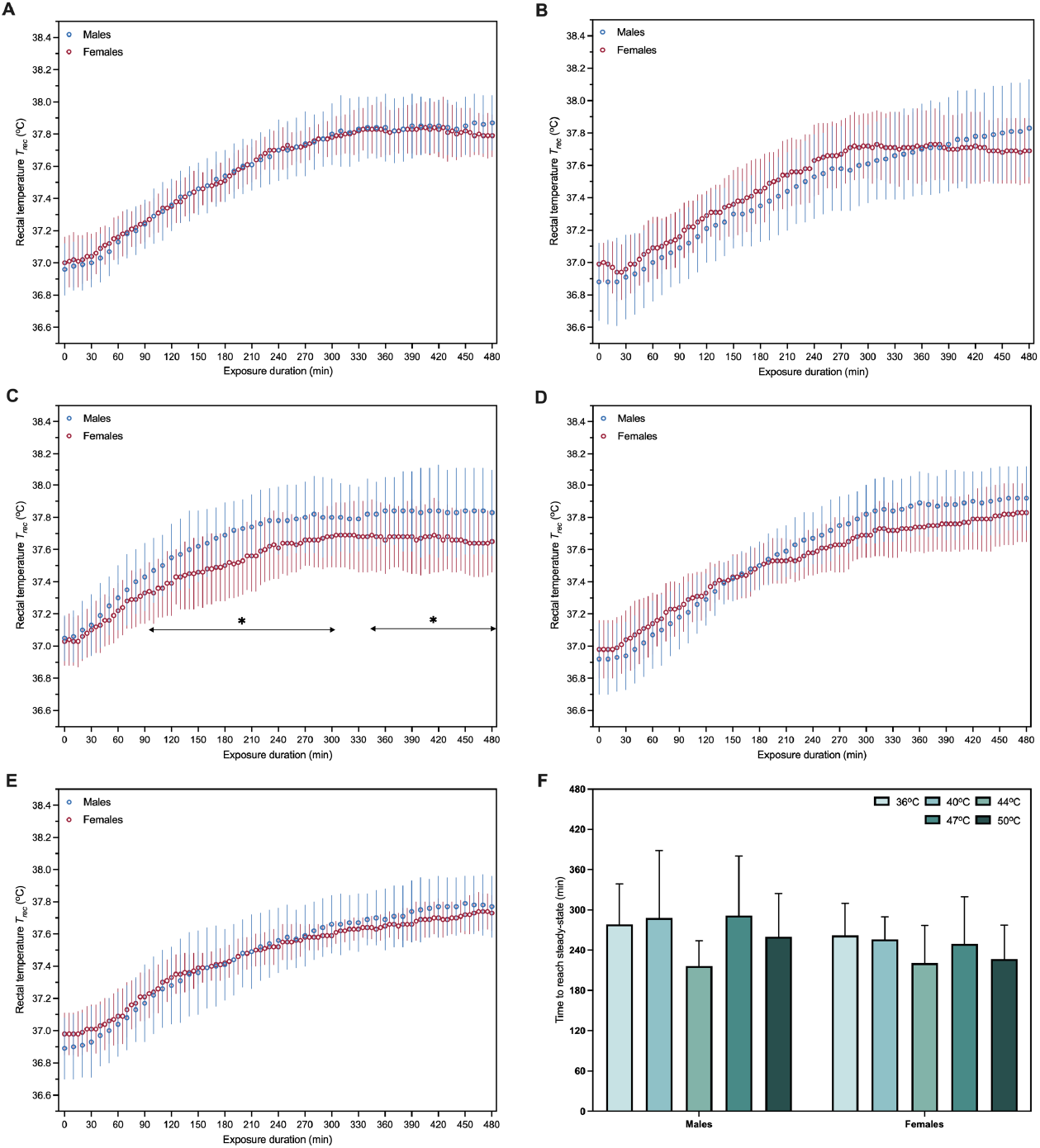
Time course of core temperature and time to reach steady-state (*T*_*rec*_ rise rate ≤ 0.10 °C/h) under five different environmental conditions. A, 36°C/74.5% RH; B, 40°C/55.0% RH; C, 44°C/29.2% RH; D, 47°C/35.6% RH; E, 50°C/24.5% RH; F, time to reach steady-state for both sexes (males: n=20; females: n=16). Data were analyzed by one-way mixed model ANOVA. *, *p* < 0.05.

Skin temperature (*T*_*sk*_) increased by the end of the 8-hour prolonged heat exposure across all five experimental conditions (all *p* < 0.01). Significant sex differences in the mean skin temperature were observed at the 40°C/55.0% RH, 44°C/29.2% RH, 47°C/35.6% RH and 50°C/24.5% RH conditions. Specifically, males exhibited higher post-exposure *T*_*sk*_ values than females at 40°C/55.0% RH, 44°C/29.2%, while females had higher post-exposure *T*_*sk*_ than males at 47°C/35.6% RH and 50°C/24.5%RH (interaction: all *p* < 0.05).

## DISCUSSION

Maintaining strict humidity control during physiological trials is crucial for accurately determining the critical environmental limit (or uncompensability limit). If the actual relative humidity exceeds the target value but remains undetected, it could lead to an underestimation of the uncompensability limit, and vice versa. Theoretically, even a 5% fluctuation in RH can significantly alter the wet-bulb temperature (*T*_*W*_). For instance, at 40°C, increasing RH from 45% to 50% raises *T*_*W*_ by 1.1°C, while at 50°C, increasing RH from 15% to 20% results in a 2.4°C rise in *T*_*W*_. The precise environmental control in our study ensures accurate assessments of environmental compensability and provides reliable validation for the predictive methods being evaluated (i.e., the CTIP and biophysical modeling). Additionally, when determining critical environmental limits, environmental temperature and relative humidity at a minimum of four heights (in our study, we used 0.1 m, 0.6 m, 1.1 m, and 1.7 m) inside the test chamber should be continuously monitored (ISO7730, 2005) and adjusted throughout the prolonged exposure to ensure the strict maintenance of the target wet-bulb temperature.

The observed metabolic rates in our study were comparable to MiniAct (minimal activity, average metabolic rate: 1.34-1.51 METs) measured in Vecellio et al.’s study (2022), as well as that used for biophysical model simulations (i.e., 1.5 METs) by Vanos et al. (2023). Additionally, we observed a 15.4% to 20.3% increase in participants’ metabolic rate after the 8-hour heat exposure, which aligns well with existing literatures reporting a comparable metabolic rate increase (i.e., 7-23%) per degree Celsius rise in core temperature (Saxton, 1981; Cramer et al., 2022). It should be noted that the participant’s metabolic rate was assumed to be constant when employing the CTIP method and biophysical modeling to estimate critical environmental limits. This fails to replicate the actual metabolic process for both methods and thereby, introducing errors to the estimated critical environmental limits (or uncompensability limit).

Our findings highlight key discrepancies between empirical data and predictions from biophysical models and the CTIP method, revealing limitations in defining the “uncompensability limit”. Biophysical models, which rely on simplified heat exchange equations, often fail to account for the complexities of human thermoregulation, behavioral adaptations, and sex differences (see Figure 3). For instance, adaptive behaviors such as periodic sweat drying, which enhances evaporative cooling, play a vital role in mitigating heat stress, particularly in high-humidity environments (Piantadosi, 2023). Our study found that participants were able to endure all five environmental conditions for prolonged durations, in contrast to model predictions that suggested a rapid onset of heatstroke. Notably, the biophysical model estimated that participants could reach a core temperature (*T*_*core*_) of 43°C within 6 hours under the 47°C with 35.6% RH and 50°C with 24.5% RH conditions. However, the actual physiological responses demonstrated greater tolerance than predicted, highlighting the limitations of the model in capturing human adaptive behaviors and thermoregulatory responses. A significant limitation of biophysical models is their lack of real-world validation under survival threshold conditions. To the best of our knowledge, models such as those used by Vanos et al. (2023) have not been tested in the extreme environments examined in our study. This absence of empirical validation likely contributes to the substantial overestimation of heat stress and underestimation of human survivability observed in their predictions.

**Figure 3.**
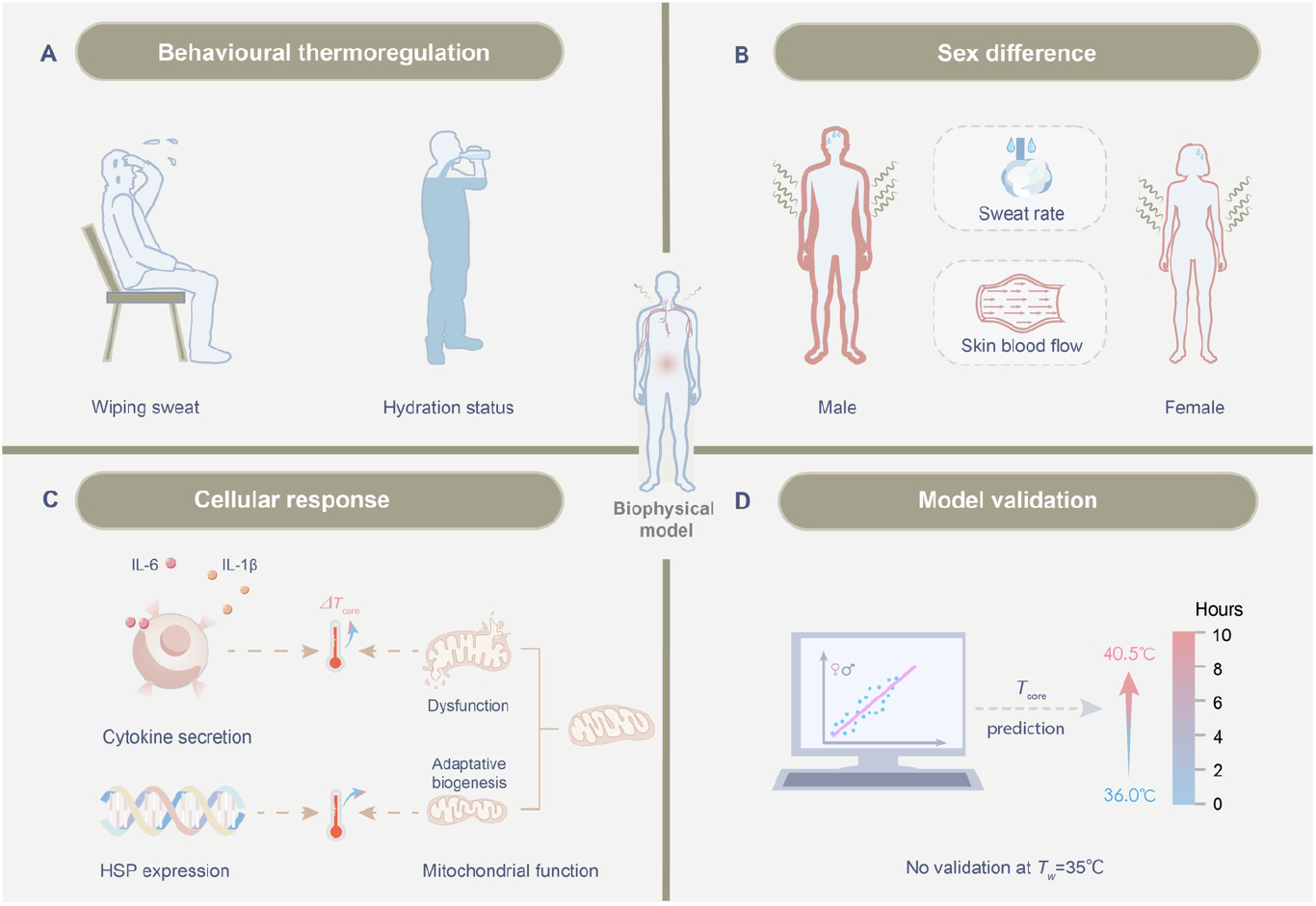
Critical gaps in biophysical models in predicting human responses to extreme heat stress. (A) Lack of consideration for behavioral thermoregulation strategies, such as hydration status and periodic skin drying (Tansey and Johnson, 2015). (B) Neglect of sex differences, including variations in sweating response, body physique, and cardiovascular capacity (Kaciuba-Uscilko and Grucza, 2001). (C) Omission of cellular response mechanisms, including mitochondrial function, cytokine secretion (e.g., IL-6, IL-1β), and heat shock protein (HSP) expression (Leon et al., 1998). These responses can significantly influence the rate and extent of core temperature (*T*_*core*_) rise. Pro-inflammatory cytokines may increase metabolic heat production and impair heat dissipation, while HSPs stabilize proteins and enhance metabolic efficiency (Fields, 2001). Additionally, adaptive mitochondrial and vascular changes improve heat tolerance over time. (D) Insufficient model validation under extreme conditions, with no testing at *T*_W_=33-35°C or during prolonged extreme heat exposures.

Most models overlook sex differences in thermoregulation. Our results showed that females exhibited slower core temperature rises than males (Figure 2C), likely due to variations in body composition, metabolic heat production, and vascular adaptations (Piantadosi, 2023). These differences suggest that females may have greater heat tolerance during prolonged uncompensable exposure. However, treating humans as a homogenous group, as most biophysical models do, leads to inaccuracies in predicting heat tolerance for mixed-sex populations. Incorporating sex-specific data into these heat tolerance models is therefore essential to improve their accuracy and applicability.

The core temperature inflection point (CTIP) method also has notable shortcomings. This method assumes that a transient rise in core temperature over a short timeframe (5 minutes) indicates a loss of heat balance, thereby signaling uncompensable heat stress and often predicting heatstroke at lower wet-bulb temperatures (e.g., critical *T*_*W*_ averaging 30.55±0.98°C) (Vecellio et al., 2022). However, such short-duration fluctuations more likely reflect a delay in thermoeffector responses rather than true heat imbalance (Meade et al., 2024; Wang, 2025). Our prolonged 8-hour trials at the three chosen conditions (36°C/74.5% RH, 40°C/55.0% RH, and 44°C/29.2% RH) demonstrated that all these conditions remain compensable in both sexes with full hydration (see Figure 2F), directly contradicting the CTIP’s reported critical wet-bulb temperatures and highlighting its limitation in real-world conditions. Additionally, our results indicate that adaptive mechanisms, such as enhanced sweating efficiency, fluid intake, and increased skin blood flow, can effectively slow the core temperature rise. While not the primary focus of our study, previous research supports the role of these adaptive mechanisms in delaying heatstroke onset and mitigating heat stress risks (Wang et al., 2024). Taken together, the CTIP method fails to account for the delayed effects of thermoregulation during prolonged exposure, potentially underestimating human survivability. Future studies should incorporate real-world factors, such as hydration status, behavioral adaptations, and physiological variability, to refine the definition of uncompensable thresholds and improve predictive accuracy of human heat tolerance.

## FUTURE PERSPECTIVES

Advancing heat tolerance research necessitates addressing current limitations by incorporating empirical data from human survival studies and capturing the full spectrum of physiological and behavioral responses to prolonged heat stress. Real-world survival mechanisms such as periodic sweat drying with tissues or towels and fanning with hands, should be incorporated into models to enhance their accuracy. Furthermore, accounting for physiological differences across sexes and populations is essential to ensure applicability across diverse scenarios and prevent overgeneralization. Refining biophysical models with these considerations will significantly improve their predictive power for human heat tolerance under extreme conditions. While the CTIP and biophysical modeling methods may offer rapid estimations, it lacks the precision and reliability of prolonged physiological exposure trials in defining uncompensable stress and human survival thresholds. Traditional prolonged physiological trials remain the gold standard for accurately assessing human resilience to extreme heat. Future physiological research should prioritize prolonged heat exposure trials to better replicate real-world conditions. Such studies would enable a deeper understanding of adaptive mechanisms, including enhanced sweating efficiency, fluid intake management, and vascular responses, which may not be evident in shorter trials (Wang, 2025). Beyond acute heat tolerance, it is also critical to investigate the chronic health impacts of prolonged heat exposure on cardiovascular, renal, and cognitive systems. Addressing these long-term risks is essential for developing effective public health interventions and climate adaptation strategies in response to escalating climate change-driven heatwaves. By tackling research priorities, future research will provide the comprehensive insights needed to mitigate health risks and enhance human resilience to extreme heat.

## LIMITATIONS

Our study did not aim to define uncompensable heat stress limits, and thus such critical environmental limits remain undetermined in this study. While our focus was validating the Core Temperature Inflection Point (CTIP) method and biophysical modeling, future research is warranted to establish precise uncompensability limits. Prolonged heat exposure trials across a range of wet-bulb temperatures could better elucidate human heat tolerance, survivability, and adaptability during extreme heatwaves, enhancing the applicability of these methods to real-world scenarios.

## CONCLUSIONS

This study demonstrates that all five tested conditions (36°C/74.5% RH, 40°C/55.0% RH, 44°C/29.2% RH, 47°C/35.6% RH, 50°C/24.5% RH) were compensable over 8 hours, with participants maintaining stable core temperatures under extreme heat stress. Contrary to predictions, both the Core Temperature Inflection Point (CTIP) method and biophysical modeling significantly underestimated the critical environmental limits (CELs), suggesting that these approaches may not accurately reflect human heat tolerance during prolonged exposure. These findings demonstrate a key gap between model predictions and observed responses, particularly for real-world durations (e.g., 8-hour exposures), whereas most methods rely on shorter (1-3 hour) trials. Our results emphasize the need to refine these methods to account for extended heat exposure, enhancing their reliability for occupational and public health applications. Future studies should investigate higher wet-bulb temperatures or longer exposure durations to more accurately define the true uncompensability threshold (or CELs) across diverse populations.

## Data availability statement

The data supporting the present findings are provided in the Supplemental Material.

## Acknowledgements

This research was financially supported by the National Excellent Young Scientist Program (grant number: 6119924022, to FW). We would also like to express our gratitude to all participants who took part in the study for their invaluable contributions to this research.

## Author contributions

All authors from this study meet the International Committee of Medical Journal Editors (ICMJE) criteria for authorship, and all those who qualify are listed. All authors had full access to and accept responsibility for the data presented in the study. FW conceived the research question and acquired funding. FW designed the trial. FW, TL, HW, YX, and HX collected data. FW, HW, TL and HX processed data. TL performed statistical analysis. FW, TL and YX created the data visualizations. FW drafted the manuscript. All authors revised and approved the final version of manuscript.

