## Supplemental Material for "Critical Environmental Limits: Assessing the Limitations of Core Temperature Inflection Point (CTIP) and Biophysical Modeling"

**by**

Faming Wang, Huijuan Xu, Tze-Huan Lei, Yi Xu, Haojian Wang, Lijuan Wang

This PDF file contains 20 figures and 1 table.

### Overview of the SUPPLEMENTAL MATERIAL

The SUPPLEMENTAL MATERIAL is divided into two primary sections that provide comprehensive details on the physiological responses of participants across the five environmental conditions.

The first section presents the individual core temperature responses (i.e., time course core temperature and change rates of core temperature) for all 36 participants (males: n=20 [M1 to M20], females: n=16 [F1 to F16]) under the different environmental conditions. This data provides a detailed view of how core temperature varied among individuals, offering insights into the physiological variations between participants and between sexes in response to specific temperature and humidity conditions. Each participant's core temperature data is shown separately, allowing for a clear understanding of individual variability and the broader trends observed across the five conditions.

The second section focuses on the hydration status marker, mean skin temperature, subjective thermal perceptions, and cardiovascular responses of both male and female participants. This data includes urine specific gravity (USG), the average skin temperature for each group, as well as reported perceptions of comfort and thermal sensation during the exposure to the different environmental conditions. Additionally, cardiovascular responses, such as heart rate and blood pressure, are presented for both genders. This section provides valuable context for understanding the broader physiological and psychological effects of environmental stressors, illustrating how hydration, perceptions and cardiovascular measures may vary across the five conditions.

Together, these two sections of supplemental material offer a thorough examination of both individual and group-level responses, contributing to a deeper understanding of how environmental factors impact human thermoregulatory responses.

### First section (ANALYSE FIGURES)

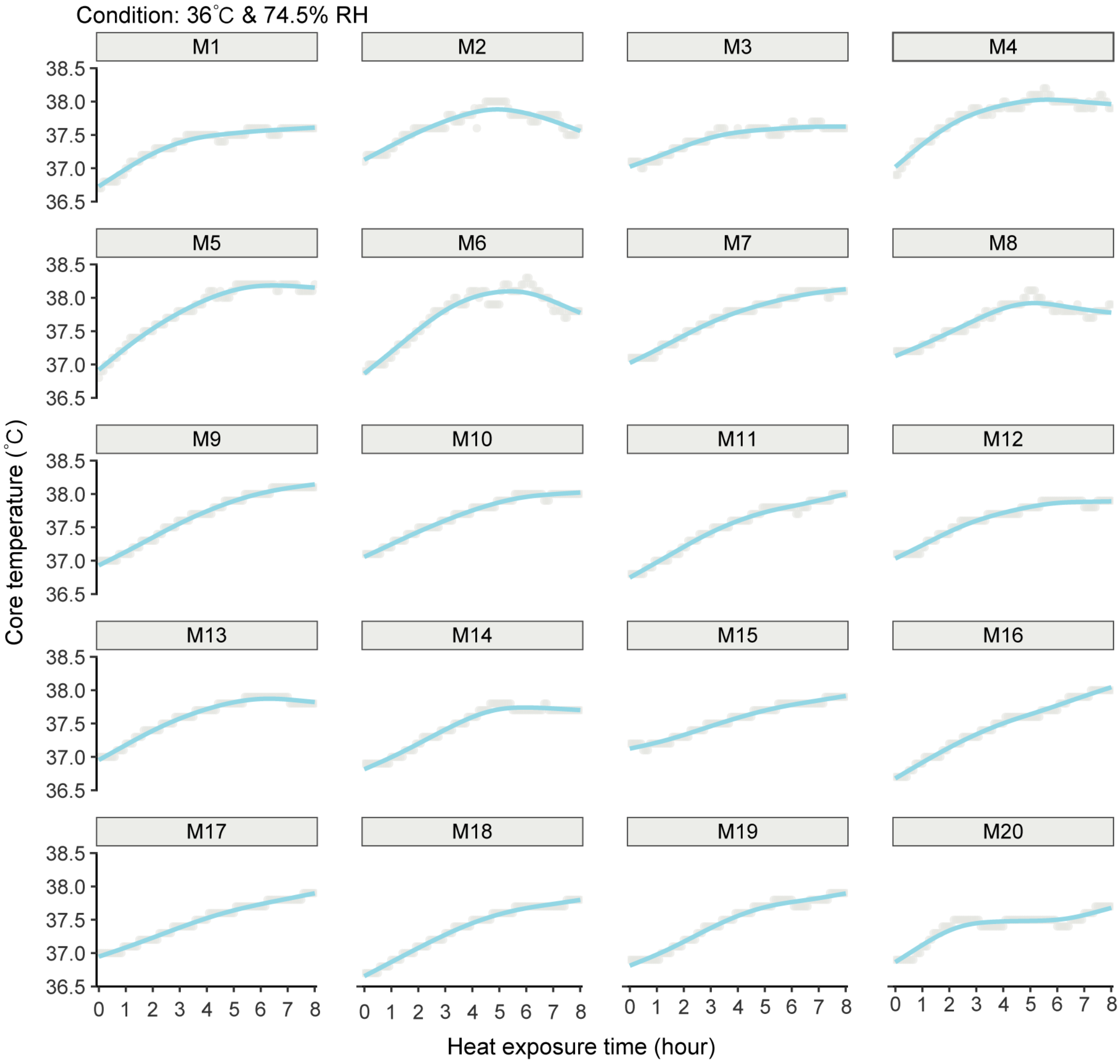


**Figure S1:** Individual core temperature responses of 20 male participants during an 8-hour heat exposure in a 36°C and 74.5% RH environment.

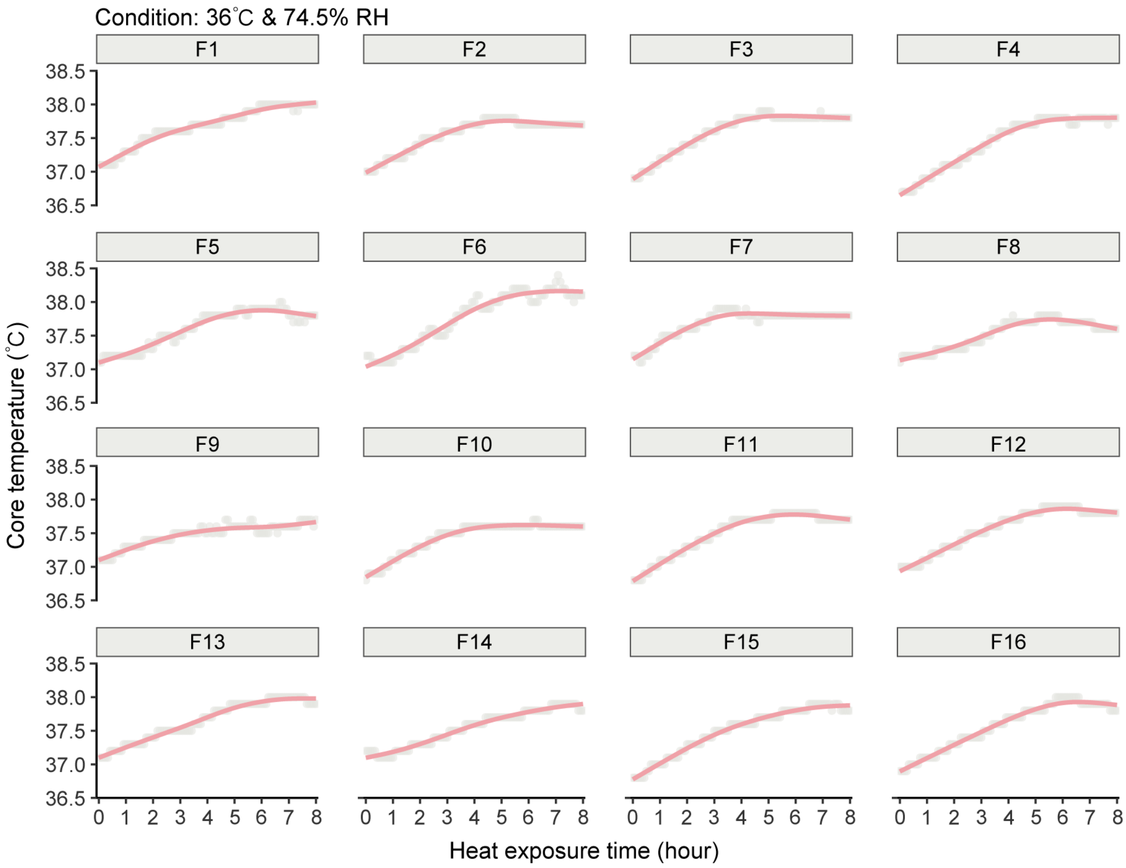


**Figure S2:** Individual core temperature responses of 16 female participants during an 8-hour heat exposure in a 36℃ and 74.5% RH environment.

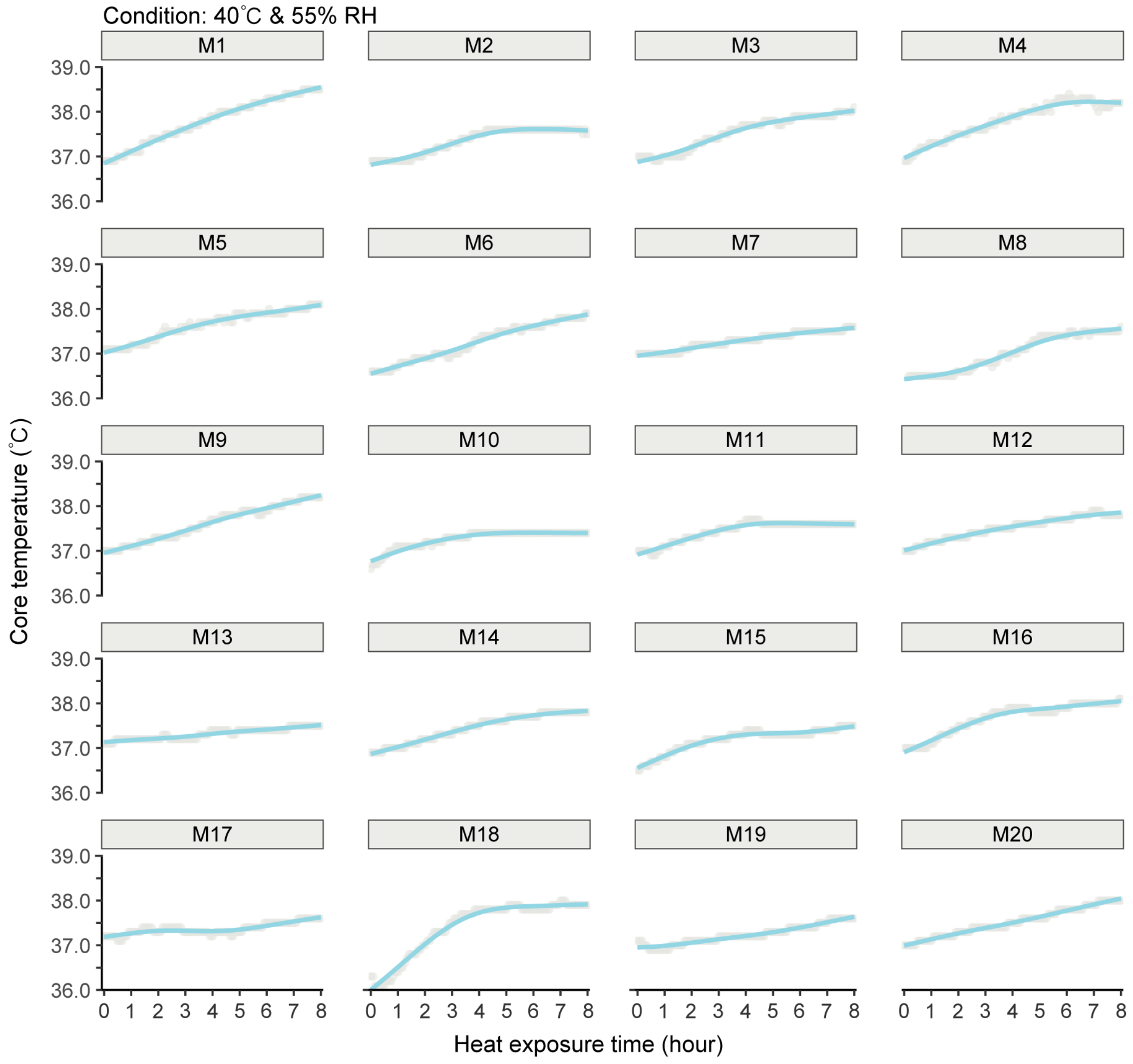


**Figure S3:** Individual core temperature responses of 20 male participants during an 8-hour heat exposure in a 40℃ and 55% RH environment.

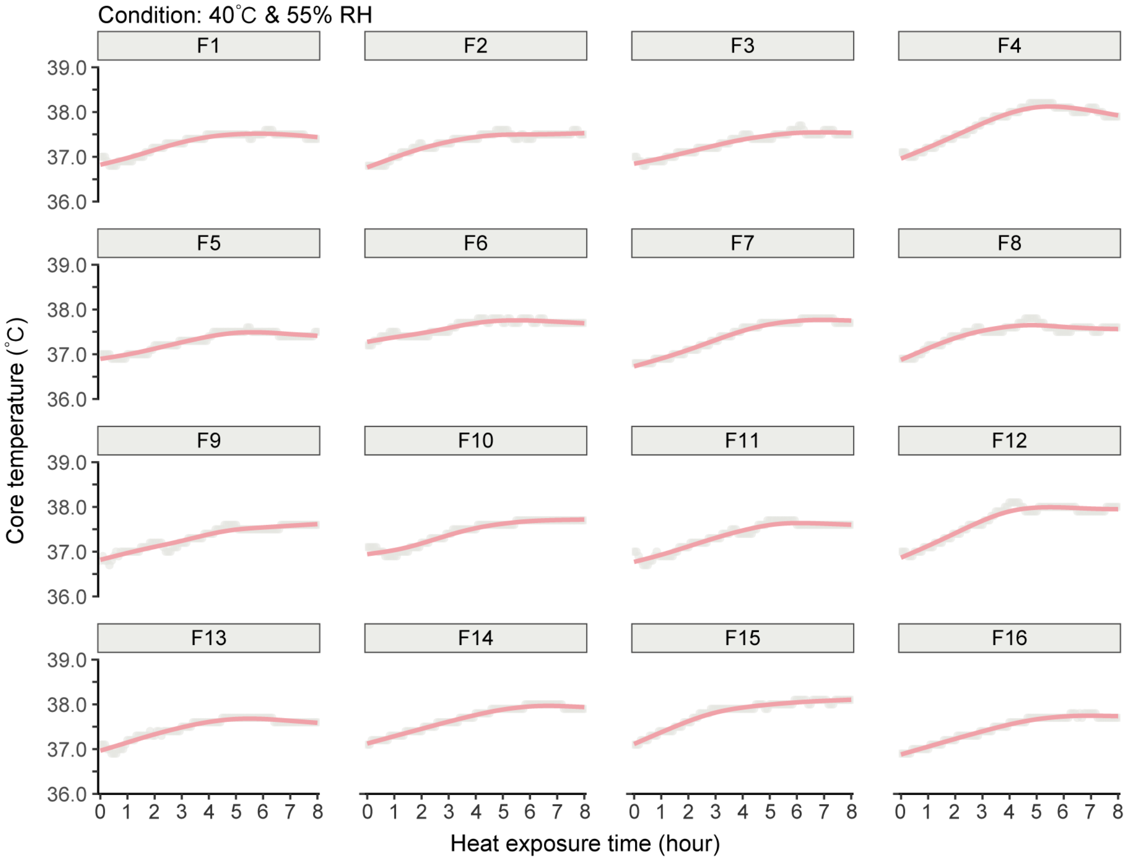


**Figure S4:** Individual core temperature responses of 16 female participants during an 8-hour heat exposure in a 40°C and 55% RH environment.

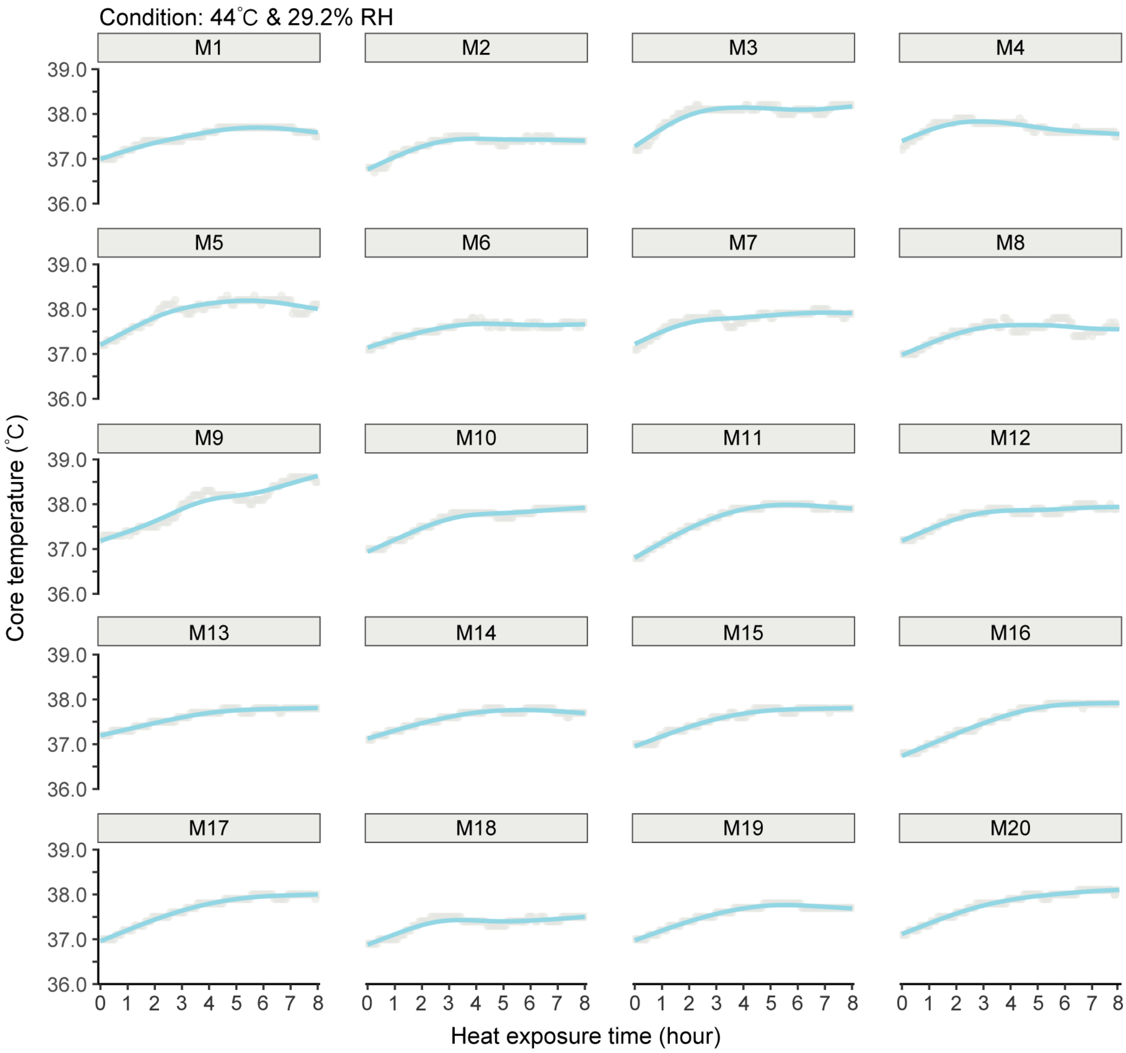


**Figure S5:** Individual core temperature responses of 20 male participants during an 8-hour heat exposure in a 44°C and 29.2% RH environment.

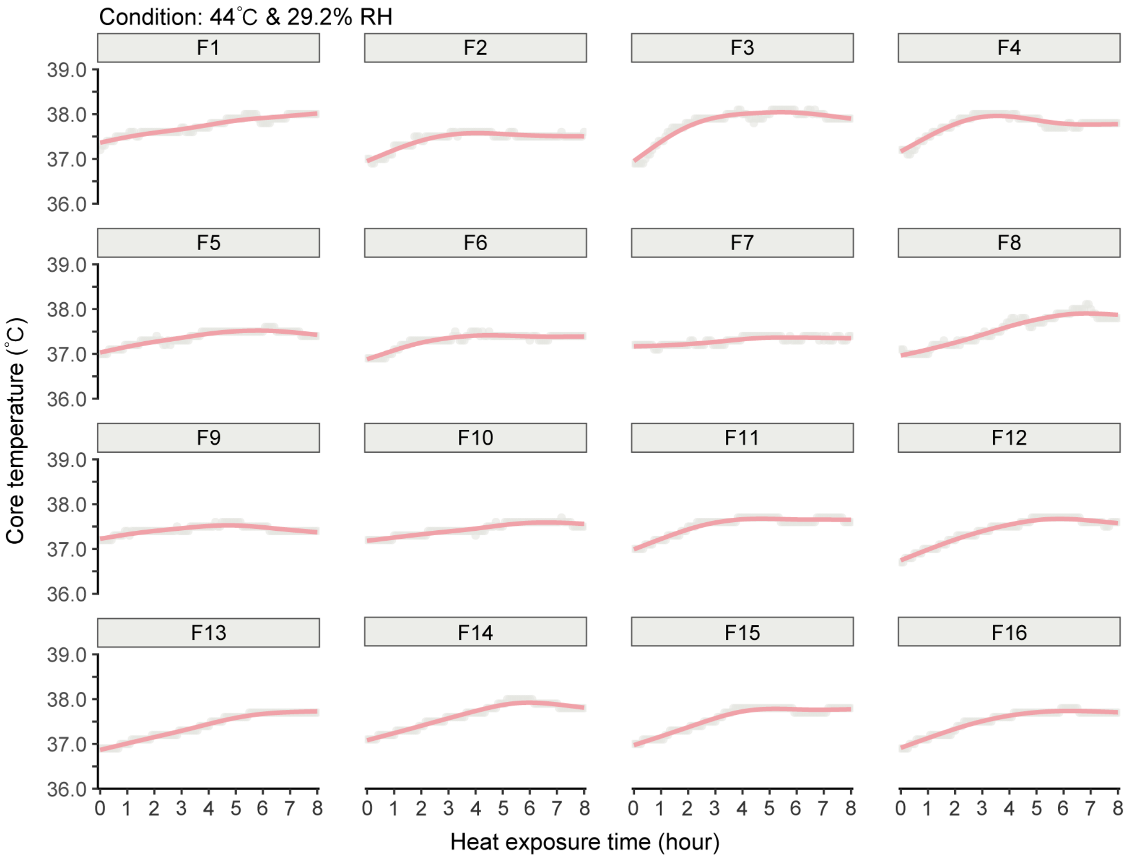


**Figure S6:** Individual core temperature responses of 16 female participants during an 8-hour heat exposure in a 44°C and 29.2% RH environment.

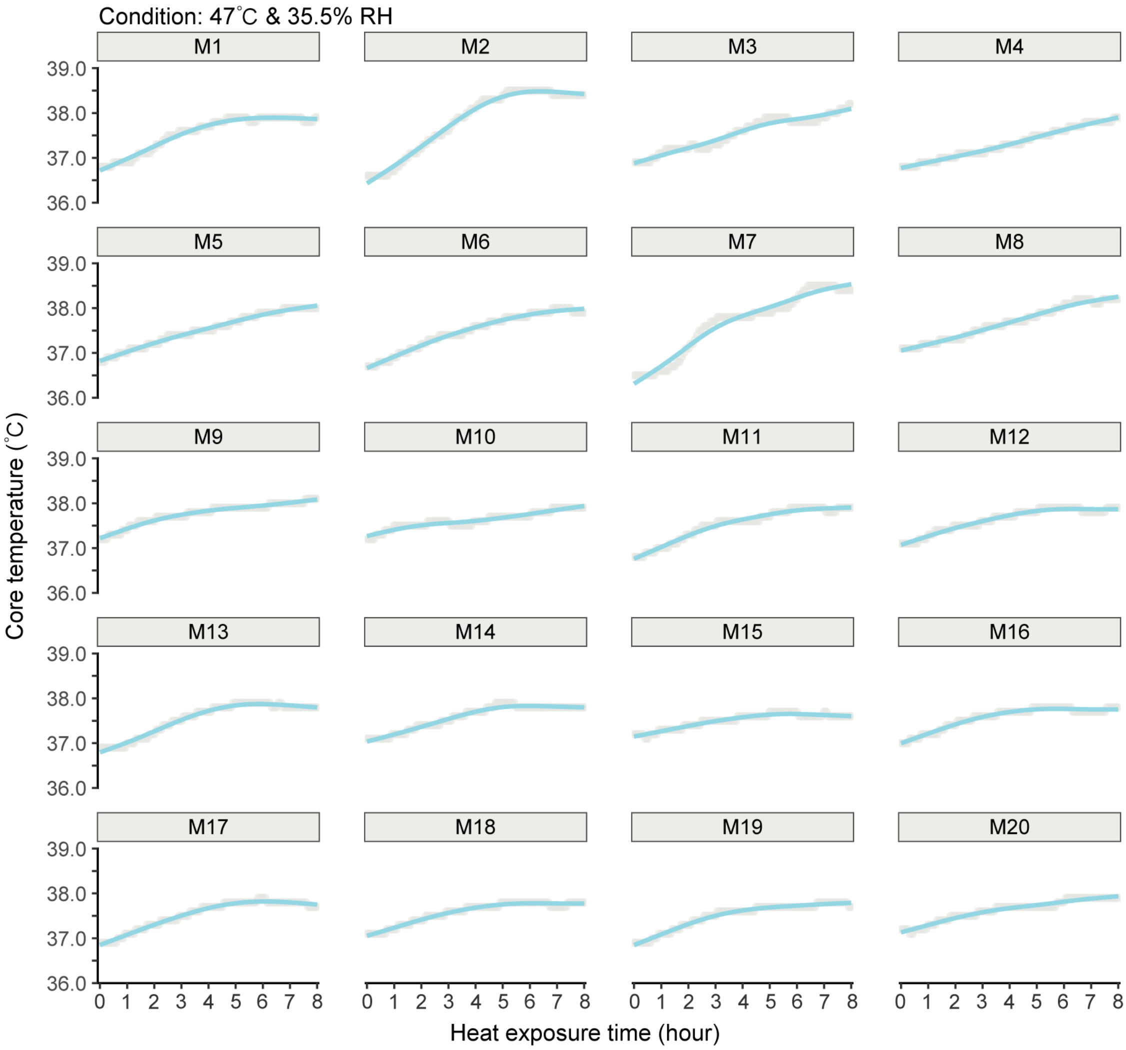


**Figure S7:** Individual core temperature responses of 20 male participants during an 8-hour heat exposure in a 47°C and 35.5% RH environment.

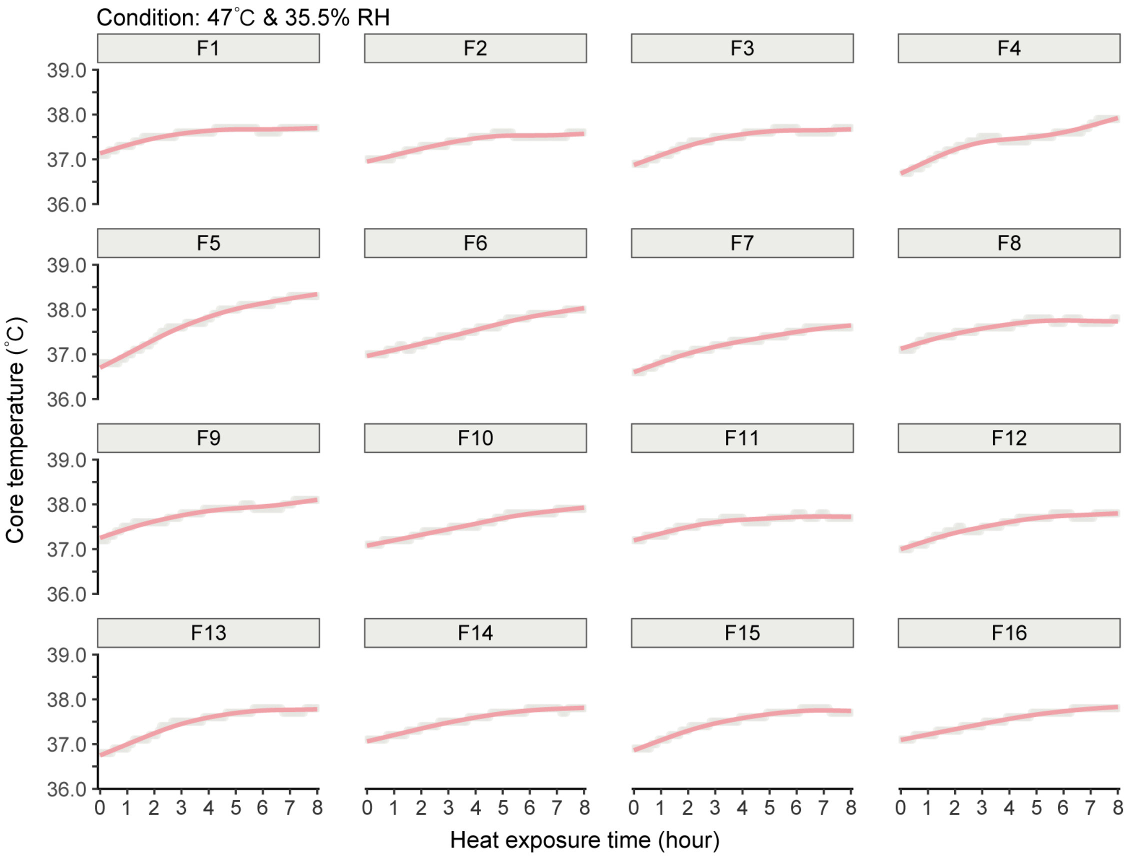


**Figure S8:** Individual core temperature responses of 16 female participants during an 8-hour heat exposure in a 47°C and 35.5% RH environment.

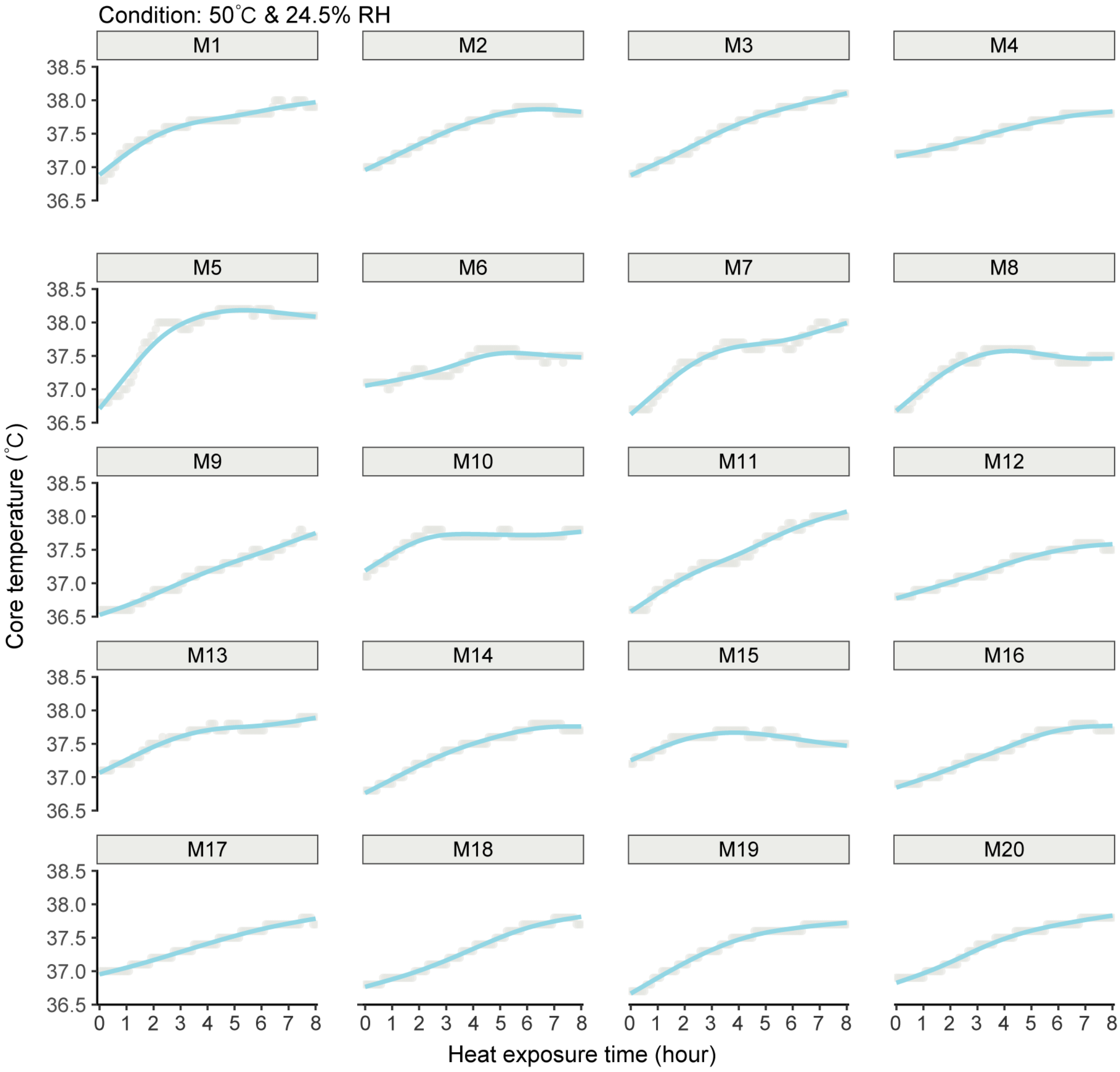


**Figure S9:** Individual core temperature responses of 20 male participants during an 8-hour heat exposure in a 50°C and 24.5% RH environment.

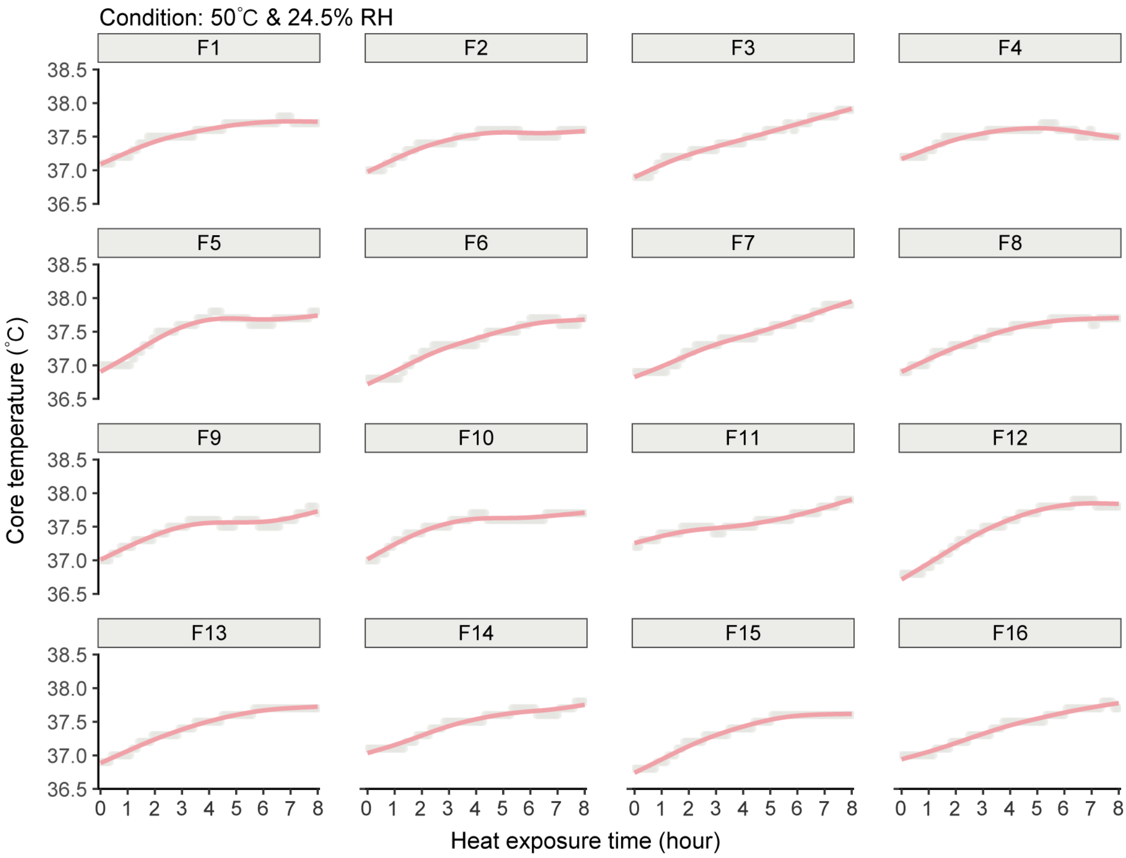


**Figure S10:** Individual core temperature responses of 16 female participants during an 8-hour heat exposure in a 50°C and 24.5% RH environment.

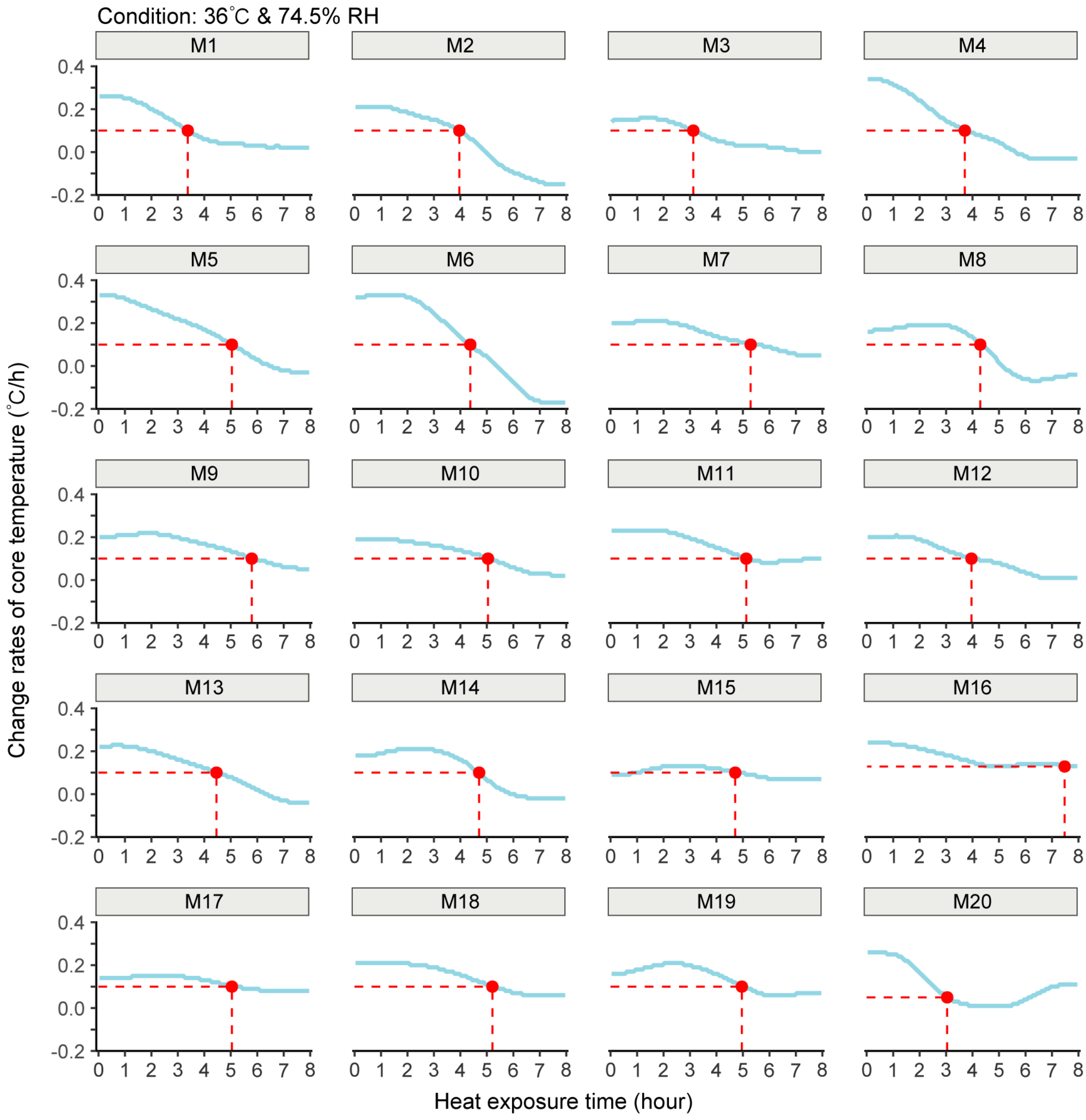


**Figure S11:** Individual rates of core temperature change for male participants (n=20) during an 8-hour heat exposure in a 36°C and 74.5% RH environment.

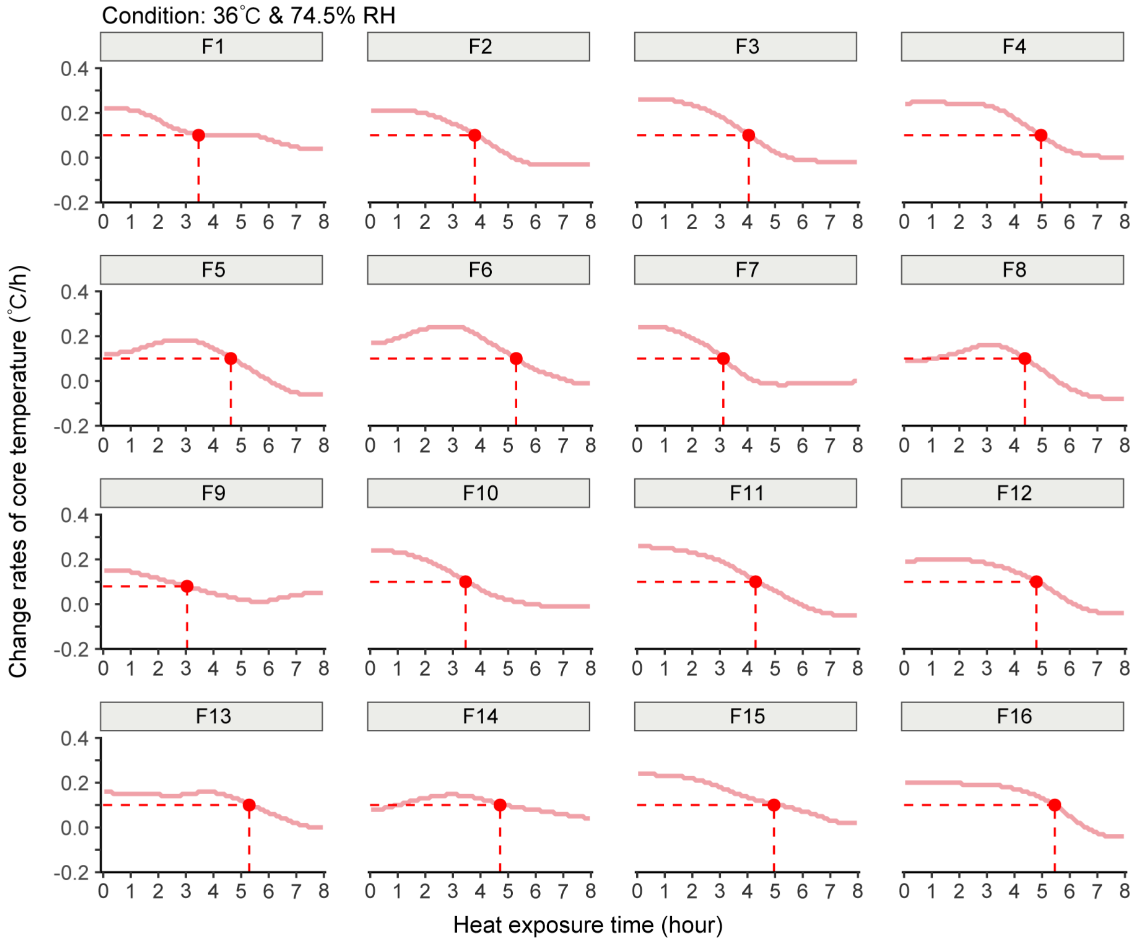


**Figure S12:** Individual rates of core temperature change for female participants (n=16) during an 8-hour heat exposure in a 36°C and 74.5% RH environment.

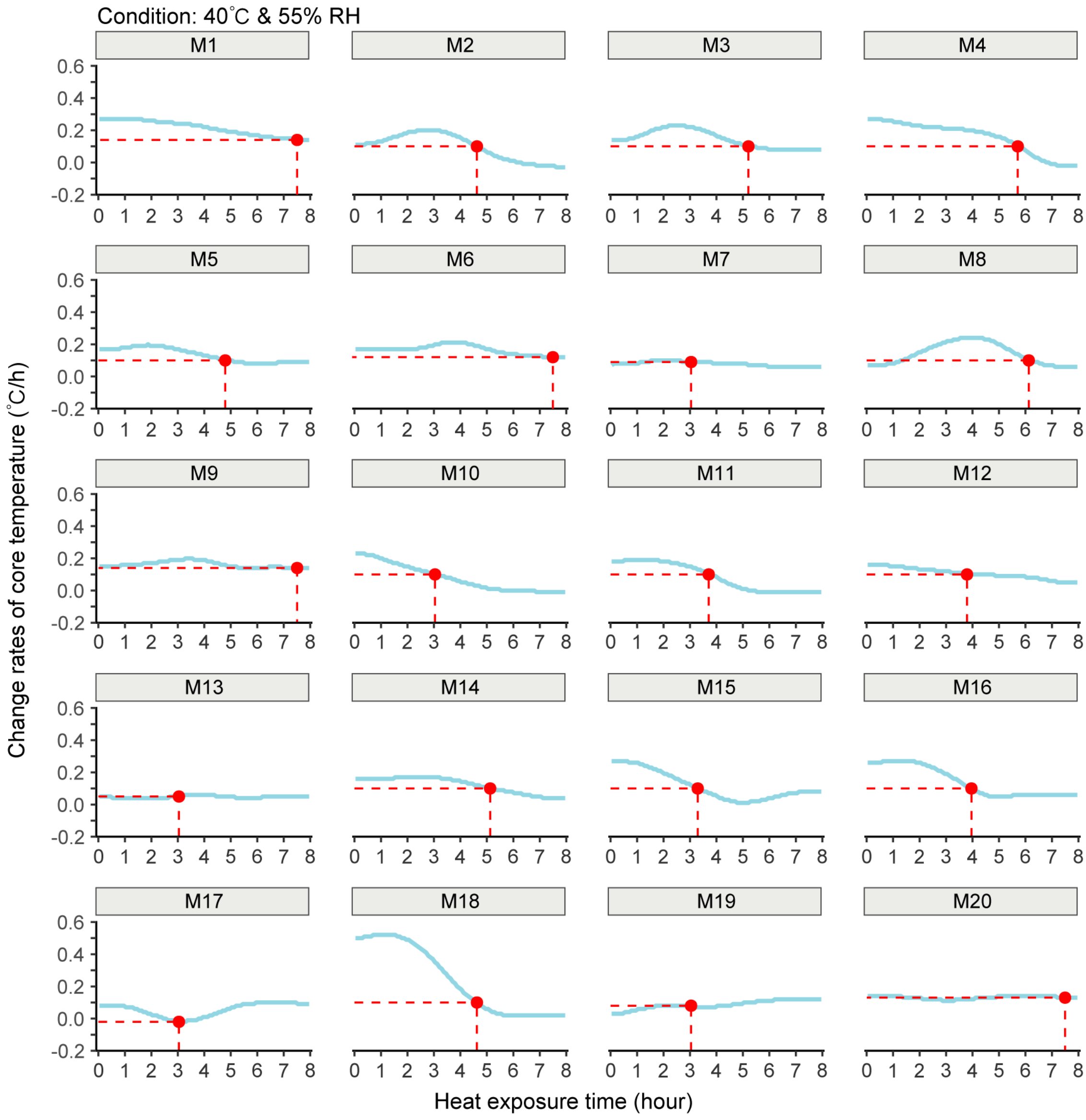


**Figure S13:** Individual rates of core temperature change for male participants (n=20) during an 8-hour heat exposure in a 40°C and 55% RH environment.

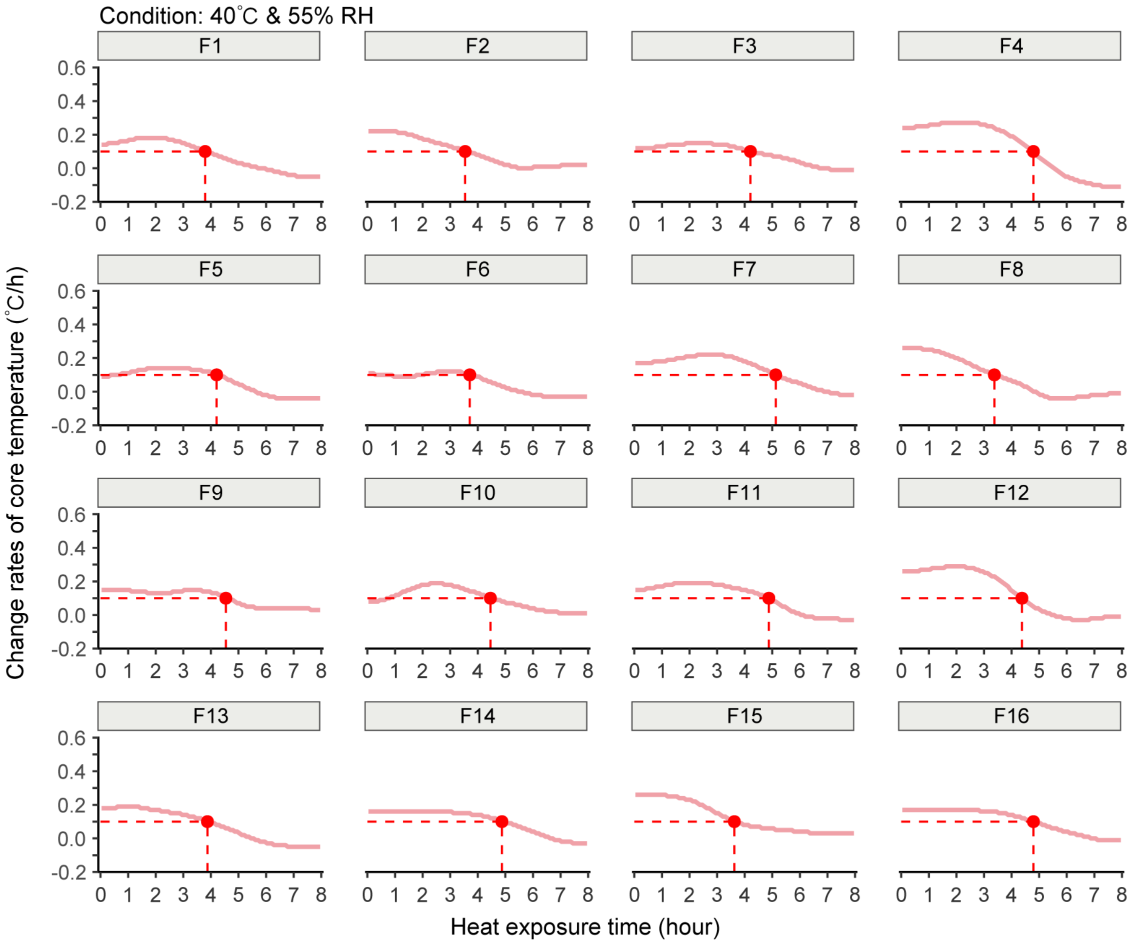


**Figure S14:** Individual rates of core temperature change for female participants (n=16) during an 8-hour heat exposure in a 40°C and 55% RH environment.

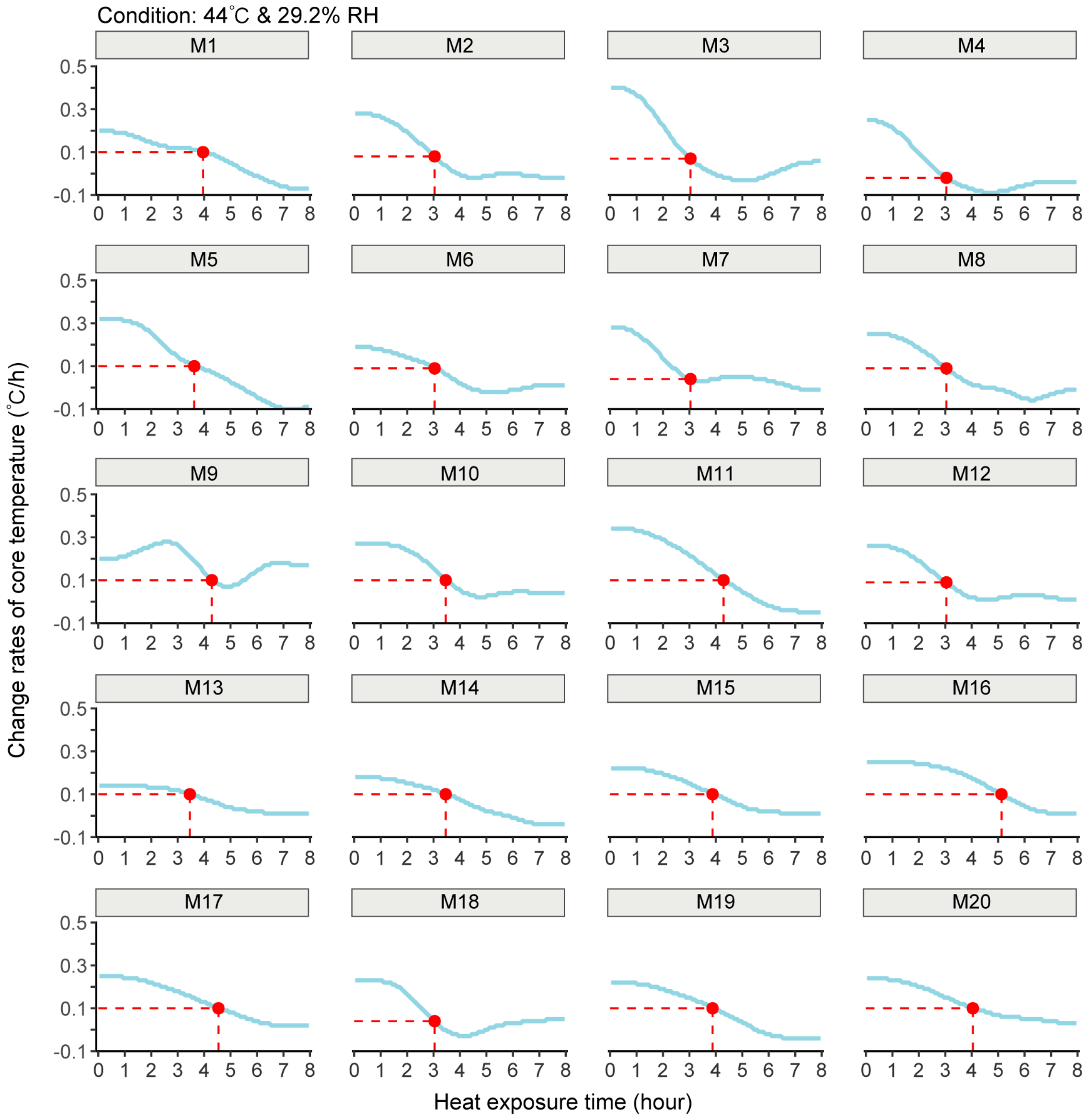


**Figure S15:** Individual rates of core temperature change for male participants (n=20) during an 8-hour heat exposure in a 44°C and 29.2% RH environment.

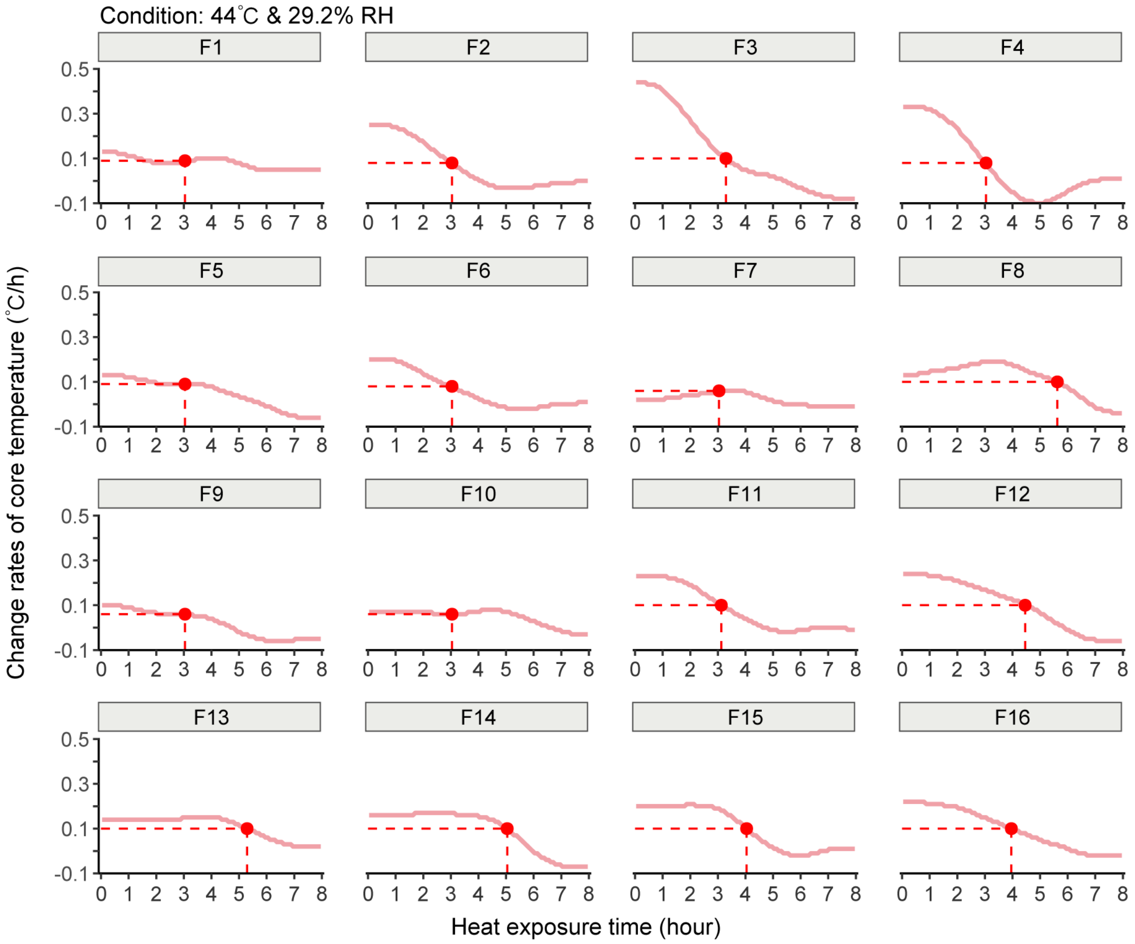


**Figure S16:** Individual rates of core temperature change for female participants (n=16) during an 8-hour heat exposure in a 44°C and 29.2% RH environment.

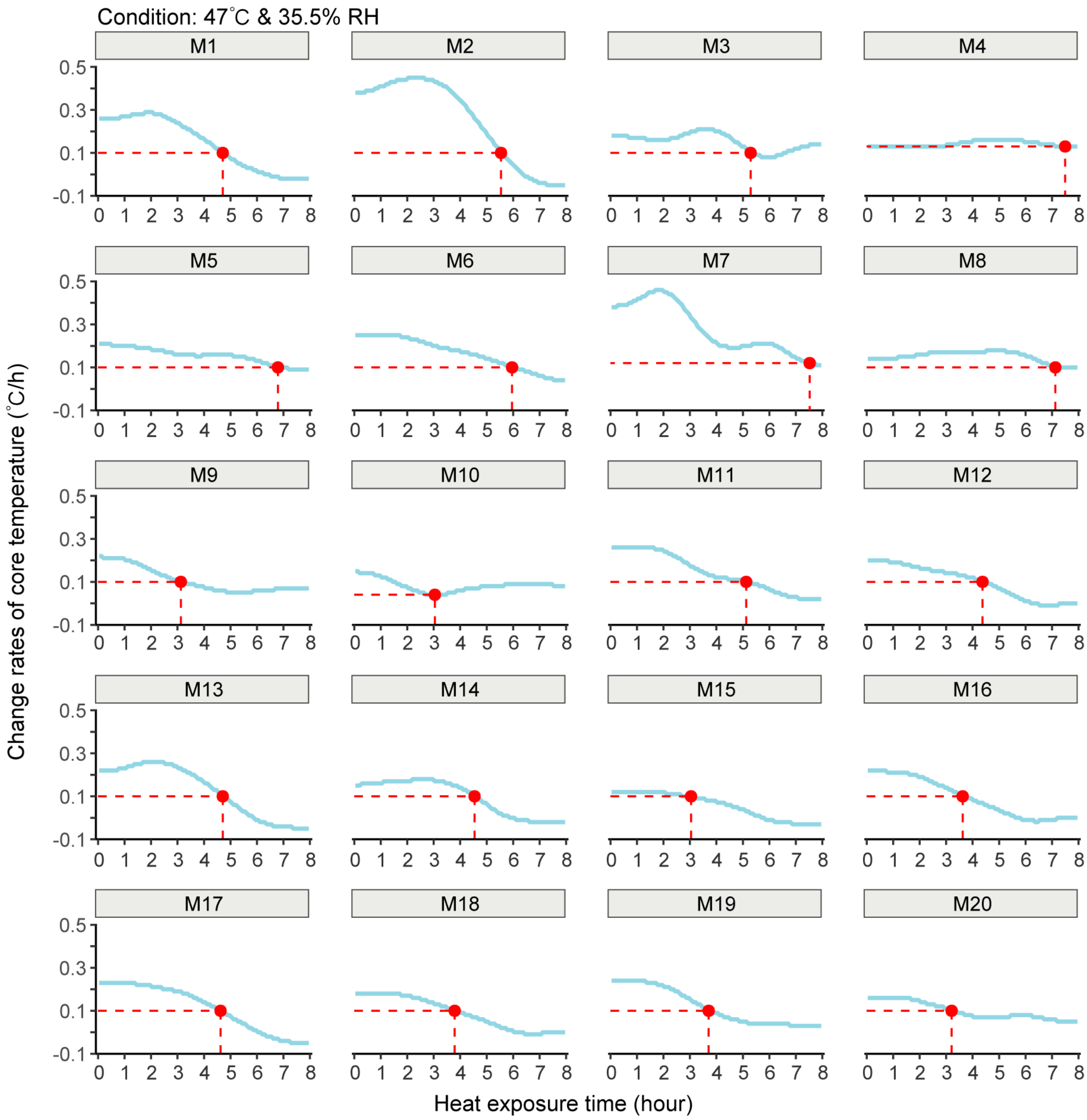


**Figure S17:** Individual rates of core temperature change for male participants (n=20) during an 8-hour heat exposure in a 47℃ and 35.6% RH environment.

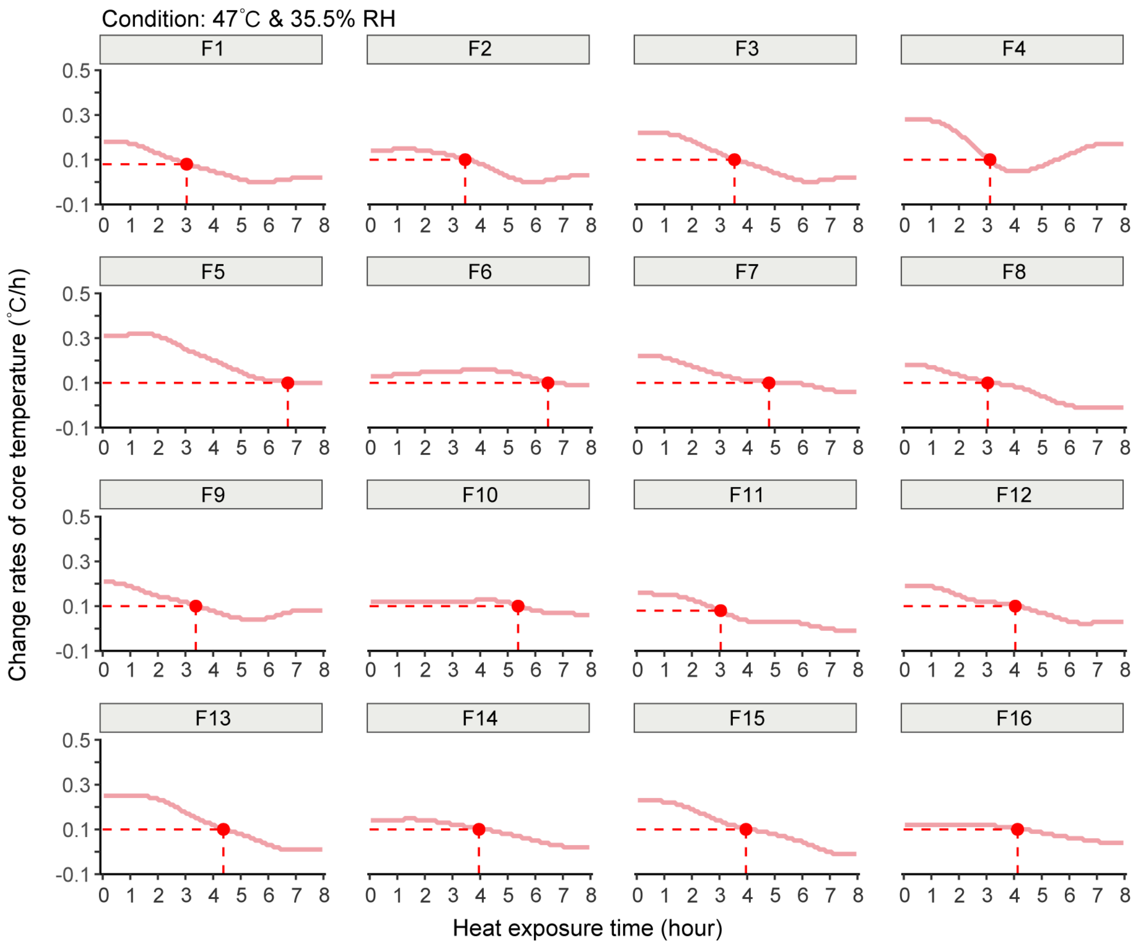


**Figure S18:** Individual rates of core temperature change for female participants (n=16) during an 8-hour heat exposure in a 47°C and 35.6% RH environment.

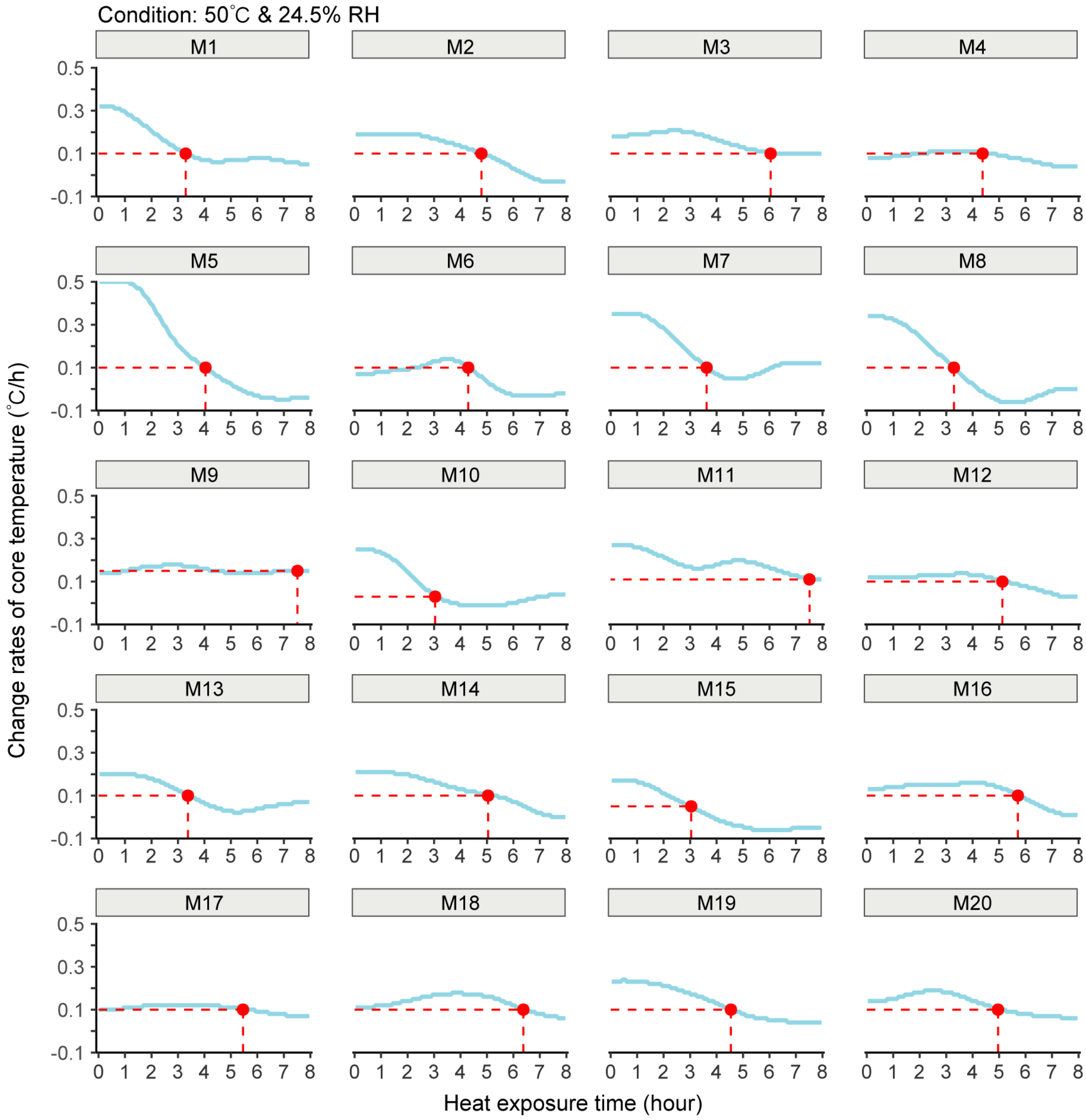


**Figure S19:** Individual rates of core temperature change for male participants (n=20) during an 8-hour heat exposure in a 50°C and 24.5% RH environment.

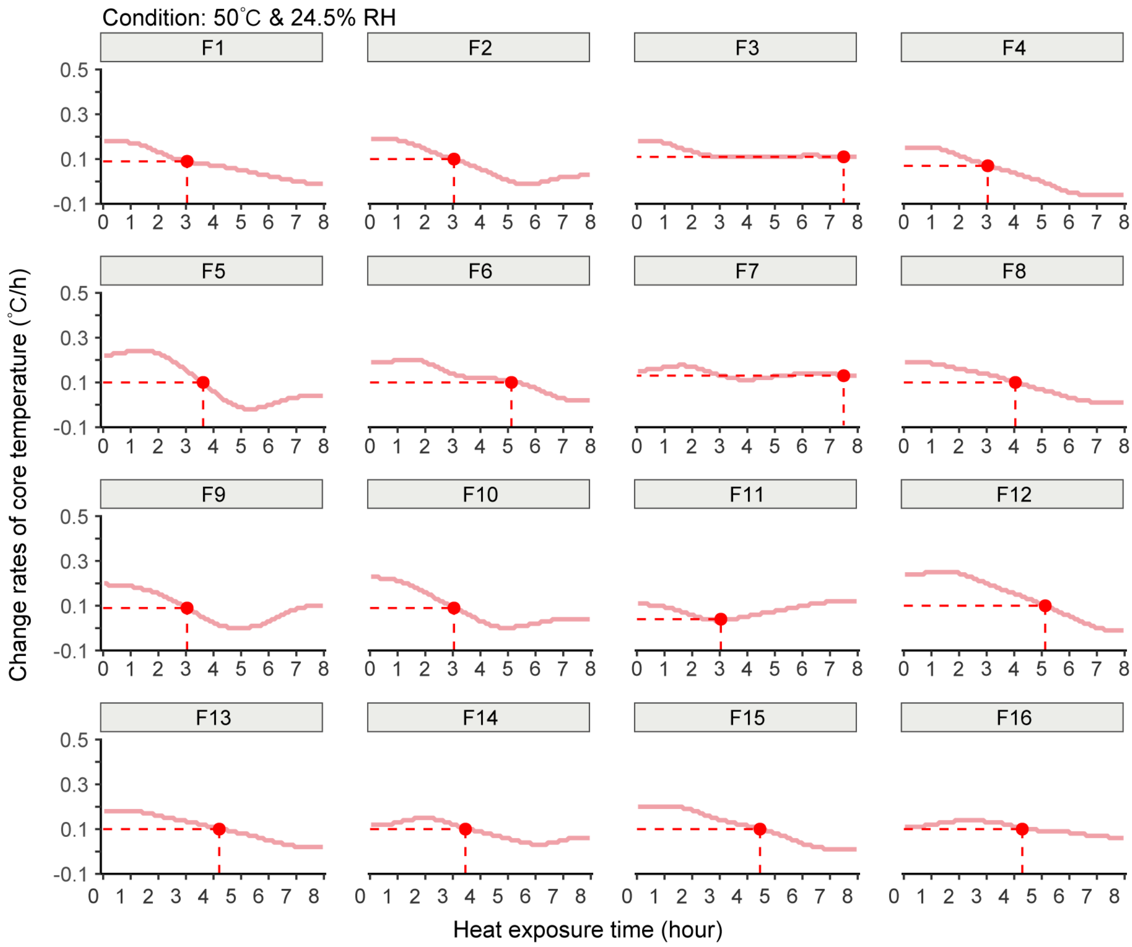


**Figure S20:** Individual rates of core temperature change for female participants (n=16) during an 8-hour heat exposure in a 50℃ and 24.5%RH environment.

### Second section

**Table S1:** Urine specific gravity (USG), mean skin temperature ($T_{sk}$), cardiovascular variables (mean arterial pressure [MAP), heart rate) and thermal perceptions (thermal sensation [TS] and thermal comfort [TC]) at pre-exposure (Pre) and at the end of the exposure (Post) for both sexes (males: n=20, females: n=16). AU, arbitrary units.

| Variables | 36°C, 74.5% RH | | | | 40°C, 55.0% RH | | | | 44°C, 29.2% RH | | | | 47°C, 35.6% RH | | | | 50°C, 24.5% RH | | | |
|---|---|---|---|---|---|---|---|---|---|---|---|---|---|---|---|---|---|---|---|---|
| | **Males** | | **Females** | | **Males** | | **Females** | | **Males** | | **Females** | | **Males** | | **Females** | | **Males** | | **Females** | |
| | Pre | Post | Pre | Post | Pre | Post | Pre | Post | Pre | Post | Pre | Post | Pre | Post | Pre | Post | Pre | Post | Pre | Post |
| **Hydration status** | | | | | | | | | | | | | | | | | | | | |
| USG (AU) | 1.016 ±0.004 | 1.013 ±0.004 | 1.018 ±0.005 | 1.012 ±0.004 | 1.018 ±0.007 | 1.018 ±0.010 | 1.012 ±0.006 | 1.007 ±0.004 | 1.017 ±0.005 | 1.015 ±0.007 | 1.019 ±0.005 | 1.012 ±0.005 | 1.013 ±0.005 | 1.020 ±0.010 | 1.018 ±0.004 | 1.020 ±0.007 | 1.014 ±0.006 | 1.020 ±0.004 | 1.016 ±0.005 | 1.020 ±0.006 |
| **Body temperature** | | | | | | | | | | | | | | | | | | | | |
| $T_{sk}$ (°C) | 33.7 ±0.4 | 37.1 ±0.4 | 33.6 ±0.3 | 36.9 ±0.2 | 34.9 ±0.3 | 37.6 ±0.4 | 33.7 ±1.4 | 37.3 ±0.1 | 33.6 ±0.4 | 37.8 ±0.5 | 33.7 ±0.2 | 37.4 ±0.3 | 32.4 ±1.0 | 38.2 ±0.3 | 32.0 ±1.0 | 38.6 ±0.4 | 33.3 ±1.6 | 38.7 ±0.3 | 32.4 ±0.4 | 39.2 ±0.5 |
| **Cardiovascular variables** | | | | | | | | | | | | | | | | | | | | |
| Heart rate (bpm) | 86 ±7 | 108 ±12 | 84 ±8 | 111 ±13 | 90 ±8 | 117 ±10 | 93 ±13 | 109 ±19 | 89 ±6 | 103 ±12 | 79 ±12 | 108 ±13 | 90 ±10 | 117 ±11 | 93 ±10 | 119 ±13 | 93 ±16 | 117 ±10 | 95 ±9 | 117 ±11 |
| MAP (mmHg) | 82 ±3 | 72 ±4 | 81 ±5 | 69 ±5 | 87 ±6 | 80 ±6 | 80 ±5 | 73 ±5 | 83 ±5 | 73 ±4 | 83 ±6 | 71 ±6 | 86 ±4 | 76 ±3 | 83 ±5 | 71 ±3 | 87 ±6 | 78 ±4 | 82 ±7 | 71 ±4 |
| **Thermal perceptions** | | | | | | | | | | | | | | | | | | | | |
| TS (AU) | 0.3 ±0.5 | 2.4 ±0.5 | 0.4 ±0.5 | 2.1 ±0.9 | 1.9 ±0.2 | 3.0 ±0 | 0.7 ±0.5 | 2.9 ±0.7 | 0.8 ±0.7 | 2.8 ±0.4 | 0.4 ±0.5 | 2.3 ±0.6 | 0.6 ±0.8 | 3.0 ±0.7 | 0.9 ±0.7 | 2.8 ±1 | 1.5 ±1.1 | 2.8 ±0.6 | 1.5 ±0.5 | 3.1 ±0.6 |
| TC (AU) | -0.1 ±0.4 | -2.3 ±0.9 | -0.1 ±0.5 | -1.8 ±1.1 | 0.3 ±0.5 | -2.0 ±0.7 | 0.3 ±0.5 | -2.9 ±0.7 | -0.4 ±0.5 | -2.6 ±0.5 | -0.3 ±0.5 | -2.1 ±0.7 | 0.6 ±0.8 | 3.0 ±0.7 | 0.9 ±0.7 | 2.8 ±1 | 0.1 ±1.1 | -1.8 ±1.2 | -0.5 ±0.5 | -2.8 ±0.6 |
